# Whole Genome Sequencing Based Analysis of Inflammation Biomarkers in the Trans-Omics for Precision Medicine (TOPMed) Consortium

**DOI:** 10.1101/2023.09.10.555215

**Authors:** Min-Zhi Jiang, Sheila M. Gaynor, Xihao Li, Eric Van Buren, Adrienne Stilp, Erin Buth, Fei Fei Wang, Regina Manansala, Stephanie M. Gogarten, Zilin Li, Linda M. Polfus, Shabnam Salimi, Joshua C. Bis, Nathan Pankratz, Lisa R. Yanek, Peter Durda, Russell P. Tracy, Stephen S. Rich, Jerome I. Rotter, Braxton D. Mitchell, Joshua P. Lewis, Bruce M. Psaty, Katherine A. Pratte, Edwin K. Silverman, Robert C. Kaplan, Christy Avery, Kari North, Rasika A. Mathias, Nauder Faraday, Honghuang Lin, Biqi Wang, April P. Carson, Arnita F. Norwood, Richard A. Gibbs, Charles Kooperberg, Jessica Lundin, Ulrike Peters, Josée Dupuis, Lifang Hou, Myriam Fornage, Emelia J. Benjamin, Alexander P. Reiner, Russell P. Bowler, Xihong Lin, Paul L. Auer, Laura M. Raffield, NHLBI Trans-Omics for Precision Medicine (TOPMed) Consortium, TOPMed Inflammation Working Group

**Author notes:** These authors contributed equally (Min-Zhi Jiang, Sheila M. Gaynor). Corresponding authors: Paul L. Auer, Ph.D. MEB M1400, Division of Biostatistics, Institute for Health and Equity, and Cancer Center, Medical College of Wisconsin, Milwaukee, WI, 53226, USA Laura M. Raffield, Ph.D. 120 Mason Farm Road, 5042 Genetic Medicine Building, Chapel Hill, NC, 27599, USA.

## Abstract

Inflammation biomarkers can provide valuable insight into the role of inflammatory processes in many diseases and conditions. Sequencing based analyses of such biomarkers can also serve as an exemplar of the genetic architecture of quantitative traits. To evaluate the biological insight, which can be provided by a multi-ancestry, whole-genome based association study, we performed a comprehensive analysis of 21 inflammation biomarkers from up to 38,465 individuals with whole-genome sequencing from the Trans-Omics for Precision Medicine (TOPMed) program. We identified 22 distinct single-variant associations across 6 traits – E-selectin, intercellular adhesion molecule 1, interleukin-6, lipoprotein-associated phospholipase A2 activity and mass, and P-selectin – that remained significant after conditioning on previously identified associations for these inflammatory biomarkers. We further expanded upon known biomarker associations by pairing the single-variant analysis with a rare variant set-based analysis that further identified 19 significant rare variant set-based associations with 5 traits. These signals were distinct from both significant single variant association signals within TOPMed and genetic signals observed in prior studies, demonstrating the complementary value of performing both single and rare variant analyses when analyzing quantitative traits. We also confirm several previously reported signals from semi-quantitative proteomics platforms. Many of these signals demonstrate the extensive allelic heterogeneity and ancestry-differentiated variant-trait associations common for inflammation biomarkers, a characteristic we hypothesize will be increasingly observed with well-powered, large-scale analyses of complex traits.

## Introduction

Chronic inflammation is a risk factor for many diseases including cardiovascular disease, asthma, cancer and diabetes (1–3). Chronic inflammation has been assessed in human cohorts using a variety of immunoassay measured biomarker traits, particularly markers of innate immune system activation such as C-reactive protein (CRP) and interleukin 6 (IL-6) (2). Though there is a strong influence of social and environmental factors, previous analyses, including genome-wide association studies (GWAS), have demonstrated an underlying genetic component to variance in these traits (4,5). Heritability of biomarkers of inflammation have been estimated, for instance, to be 25-60% (6,7) for IL-6 and 30-45% (8–12) for CRP. However, most studies have only analyzed relatively small and ancestrally homogenous (mostly European ancestry) populations and as such have not fully elucidated the genetic influence on these traits (4,13–16).

The National Heart Lung and Blood Institute’s Trans-Omics for Precision Medicine (TOPMed) initiative has now generated whole genome sequencing data on >150,000 individuals from diverse population-based cohorts enriched for heart, lung, and blood relevant disease traits. Novel ancestry-differentiated variant associations for CRP (17) (including confirmation of regulatory impacts *in vitro*) and E-selectin (18) reported in earlier TOPMed publications demonstrated the potential for genetic discovery for inflammation traits in these diverse cohorts. Thus, analysis of more biomarkers across a larger, more diverse set of samples with the addition of rare variant aggregate tests may identify additional associated individual variants and genomic regions. Here, we perform single variant and aggregate rare variant analyses across 21 inflammation-related biomarkers, some of which are in moderate to low correlation (Figure S1), assessed in TOPMed cohort studies (Table 1), including performing detailed conditional analyses to identify distinct genetic association signals. Our results both inform our understanding of inflammation trait biology and of the expected findings for sequencing-based analyses of complex traits, particularly protein quantitative biomarkers.

**Table 1.**
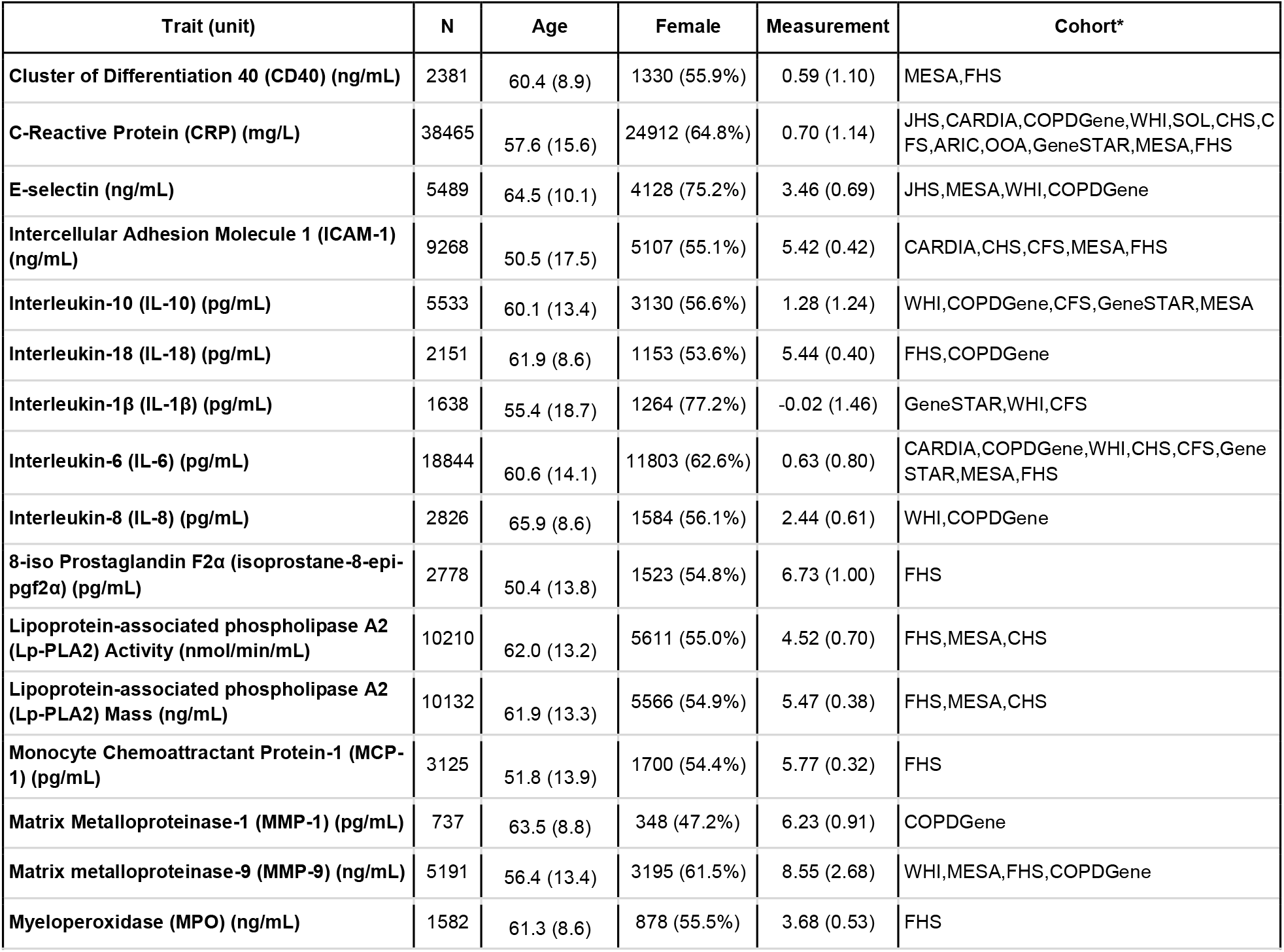
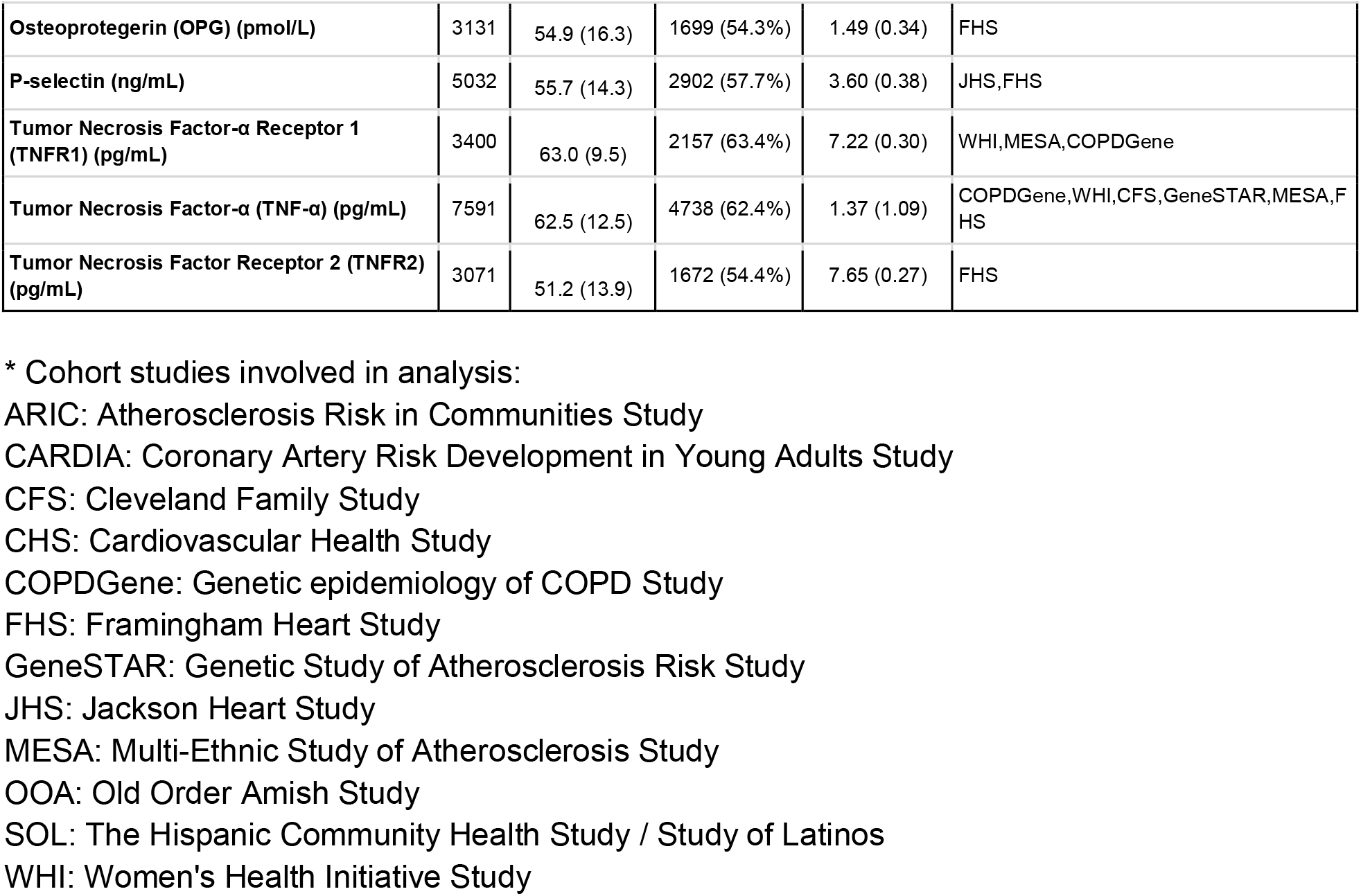
Overview of 21 inflammation-related biomarkers.

| Trait (unit) | N | Age | Female | Measurement | Cohort* |
| --- | --- | --- | --- | --- | --- |
| Cluster of Differentiation 40 (CD40) (ng/mL) | 2381 | 60.4 (8.9) | 1330 (55.9%) | 0.59 (1.10) | MESA,FHS |
| C-Reactive Protein (CRP) (mg/L) | 38465 | 57.6 (15.6) | 24912 (64.8%) | 0.70 (1.14) | JHS,CARDIA,COPDGene,WHI,SOL,CHS,CFS,ARIC,OOA,GeneSTAR,MESA,FHS |
| E-selectin (ng/mL) | 5489 | 64.5 (10.1) | 4128 (75.2%) | 3.46 (0.69) | JHS,MESA,WHI,COPDGene |
| Intercellular Adhesion Molecule 1 (ICAM-1) (ng/mL) | 9268 | 50.5 (17.5) | 5107 (55.1%) | 5.42 (0.42) | CARDIA,CHS,CFS,MESA,FHS |
| Interleukin-10 (IL-10) (pg/mL) | 5533 | 60.1 (13.4) | 3130 (56.6%) | 1.28 (1.24) | WHI,COPDGene,CFS,GeneSTAR,MESA |
| Interleukin-18 (IL-18) (pg/mL) | 2151 | 61.9 (8.6) | 1153 (53.6%) | 5.44 (0.40) | FHS,COPDGene |
| Interleukin-1β (IL-1β) (pg/mL) | 1638 | 55.4 (18.7) | 1264 (77.2%) | -0.02 (1.46) | GeneSTAR,WHI,CFS |
| Interleukin-6 (IL-6) (pg/mL) | 18844 | 60.6 (14.1) | 11803 (62.6%) | 0.63 (0.80) | CARDIA,COPDGene,WHI,CHS,CFS,GeneSTAR,MESA,FHS |
| Interleukin-8 (IL-8) (pg/mL) | 2826 | 65.9 (8.6) | 1584 (56.1%) | 2.44 (0.61) | WHI,COPDGene |
| 8-iso Prostaglandin F2α (isoprostane-8-epi-pgf2α) (pg/mL) | 2778 | 50.4 (13.8) | 1523 (54.8%) | 6.73 (1.00) | FHS |
| Lipoprotein-associated phospholipase A2 (Lp-PLA2) Activity (nmol/min/mL) | 10210 | 62.0 (13.2) | 5611 (55.0%) | 4.52 (0.70) | FHS,MESA,CHS |
| Lipoprotein-associated phospholipase A2 (Lp-PLA2) Mass (ng/mL) | 10132 | 61.9 (13.3) | 5566 (54.9%) | 5.47 (0.38) | FHS,MESA,CHS |
| Monocyte Chemoattractant Protein-1 (MCP-1) (pg/mL) | 3125 | 51.8 (13.9) | 1700 (54.4%) | 5.77 (0.32) | FHS |
| Matrix Metalloproteinase-1 (MMP-1) (pg/mL) | 737 | 63.5 (8.8) | 348 (47.2%) | 6.23 (0.91) | COPDGene |
| Matrix metalloproteinase-9 (MMP-9) (ng/mL) | 5191 | 56.4 (13.4) | 3195 (61.5%) | 8.55 (2.68) | WHI,MESA,FHS,COPDGene |
| Myeloperoxidase (MPO) (ng/mL) | 1582 | 61.3 (8.6) | 878 (55.5%) | 3.68 (0.53) | FHS |
| Osteoprotegerin (OPG) (pmol/L) | 3131 | 54.9 (16.3) | 1699 (54.3%) | 1.49 (0.34) | FHS |
| P-selectin (ng/mL) | 5032 | 55.7 (14.3) | 2902 (57.7%) | 3.60 (0.38) | JHS,FHS |
| Tumor Necrosis Factor-α Receptor 1 (TNFR1) (pg/mL) | 3400 | 63.0 (9.5) | 2157 (63.4%) | 7.22 (0.30) | WHI,MESA,COPDGene |
| Tumor Necrosis Factor-α (TNF-α) (pg/mL) | 7591 | 62.5 (12.5) | 4738 (62.4%) | 1.37 (1.09) | COPDGene,WHI,CFS,GeneSTAR,MESA,FHS |
| Tumor Necrosis Factor Receptor 2 (TNFR2) (pg/mL) | 3071 | 51.2 (13.9) | 1672 (54.4%) | 7.65 (0.27) | FHS |

\* Cohort studies involved in analysis:
ARIC: Atherosclerosis Risk in Communities Study
CARDIA: Coronary Artery Risk Development in Young Adults Study
CFS: Cleveland Family Study
CHS: Cardiovascular Health Study
COPDGene: Genetic epidemiology of COPD Study
FHS: Framingham Heart Study
GeneSTAR: Genetic Study of Atherosclerosis Risk Study
JHS: Jackson Heart Study
MESA: Multi-Ethnic Study of Atherosclerosis Study
OOA: Old Order Amish Study
SOL: The Hispanic Community Health Study / Study of Latinos
WHI: Women's Health Initiative Study

## Results

Our analyses of 21 inflammation biomarkers, generally measured by ELISA, included 12 cohorts from the TOPMed Program (Table S1); phenotype availability varied by trait (Table S3). In brief, we performed single variant analysis to identify trait-associated loci, followed by stepwise conditional analysis to identify the total number of statistically distinct signals. We also conditioned on previously associated variants to identify distinct signals not identified in prior papers. We performed genetic region and gene centric rare variant set-based analyses for each trait and likewise conditioned on previously identified signals and distinct single variant signals that remained significant when conditioned on variants from previous GWAS (as listed in Table S6).

Of the 21 traits tested, CRP, E-selectin, intercellular adhesion molecule 1 (ICAM-1), interleukin 18 (IL-18), IL-6, lipoprotein-associated phospholipase A2 (Lp-PLA2) activity and mass, monocyte chemoattractant protein-1 (MCP-1), matrix metalloproteinase-9 (MMP-9), P-selectin, and tumor necrosis factor α receptor 2 (TNFR2) had at least 1 genome-wide significant locus in single variant analyses. Across these 11 traits there were a total of 30 genome-wide significant loci (p<1.0×10^-9^ (21)) (Table S4, Figures S2-S31, and S32), for which stepwise conditional analysis revealed a total of 67 distinct signals (Table S5). After conditioning on previously identified associations (Table S6), 22 conditionally distinct variants across 8 loci remained statistically significant for 6 traits (Table S7 and Table 2, Figure 1), and 1 trait (MMP-9) had a locus not reported in the GWAS catalog (Table 2, Figure 1). In aggregate rare variant analyses, we detected 51 significant gene-centric sets associated with 6 traits (Table S10A) and 214 significant 2-kb sliding windows associated with 7 traits (Table S11A). We observed 19 significant rare variant aggregate test associations (some in overlapping or adjoining regions) after conditioning on known variants from the GWAS catalog and single-variant signals in the present analysis (Table 3), with traits P-selectin, ICAM-1, CRP, Lp-PLA2 activity and mass, all of which also had conditionally distinct single variant results (Tables S10C and S11C). If possible, we attempted to replicate distinct single variant findings using semiquantitative inflammation biomarker measures from the SomaScan or Olink platforms in independent samples (Table 4); unfortunately, additional quantitative immunoassay measurement data for replication purposes were not available, particularly in those with WGS, so replication of rare variant aggregate test signals was not possible at the time of analysis. However, we note that all distinct rare variant aggregation signals were in known regions, increasing the plausibility of their association with inflammation traits.

**Figure 1.**
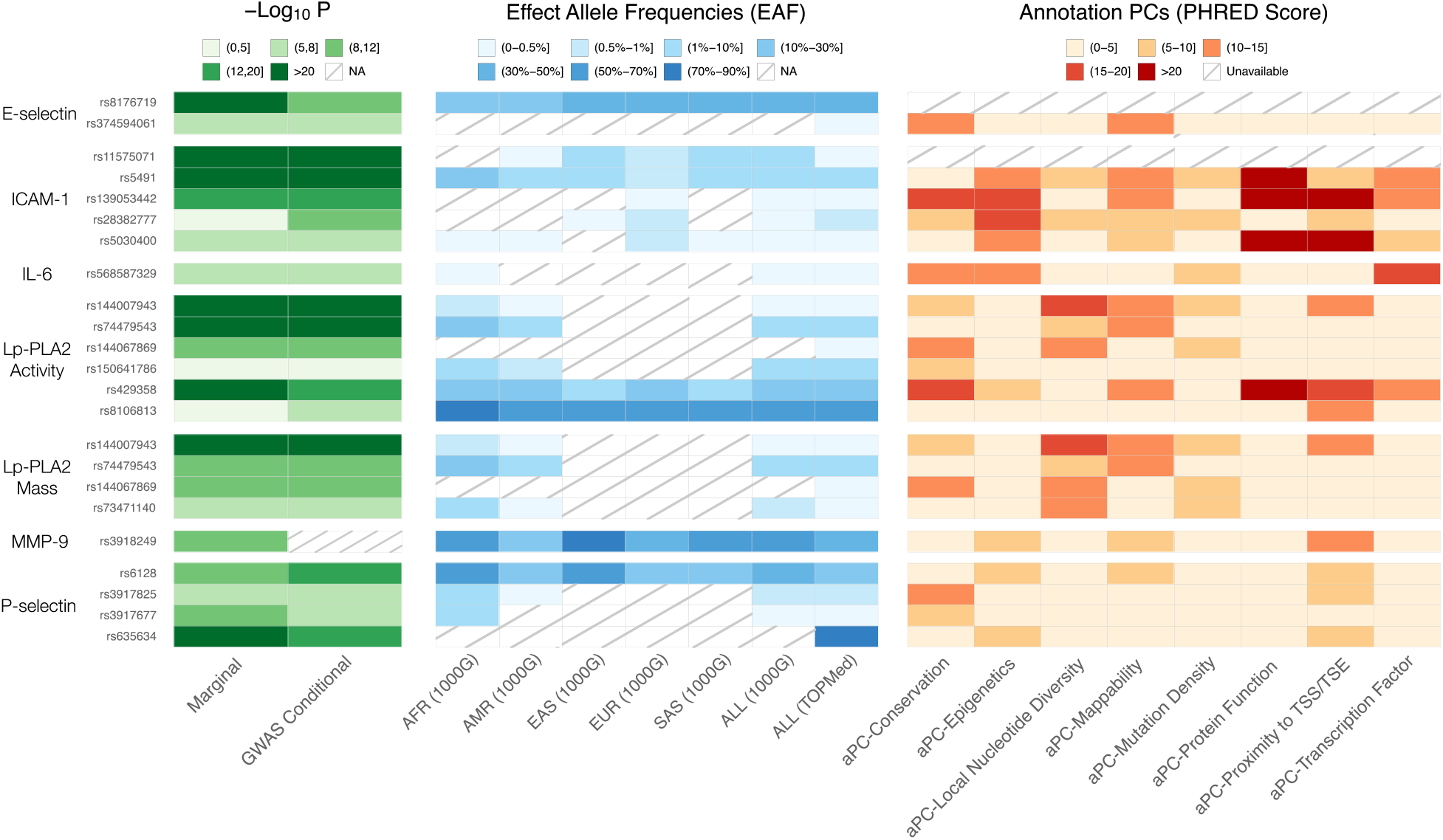
Single variant findings conditionally distinct from GWAS catalog variants. We report p-values for association for marginal and conditional results, reference population effect allele frequencies (EAF) by continental ancestry group as defined by 1000 Genomes Project (1000G) (19) – African (AFR), Admixed American (AMR), East Asian (EAS), European (EUR), South Asian (SAS), as well as all participants in 1000G (ALL) – and the overall effect allele frequency for all participants included in our TOPMed analyses, and annotation principal components (aPCs) from FAVOR (20). NA means the variant is not reported in the reference panel. We note that this information is available for all variants in close linkage disequilibrium with these lead variants in Table S8.

**Table 2.** Lead single variant signals at distinct locus (*MMP9*) and signals distinct from prior GWAS identified variants at known loci.

| Trait | Locus Name | rsID | CHR | POS (hg38) | Effect | Other | Unconditional |  |  | Conditional |  |  | EAF | Distal/Local |
| --- | --- | --- | --- | --- | --- | --- | --- | --- | --- | --- | --- | --- | --- | --- |
|  |  |  |  |  |  |  | p-value | Beta | SE | p-value | Beta | SE |  |  |
| E-selectin | *ABO* | rs8176719 | 9 | 133257521 | TC | T | 4.3E-141 | -0.24 | 0.01 | 7.7E-12 | -0.07 | 0.01 | 33.9% | Distal |
|  | *ABO* | rs374594061 | 9 | 132553865 | A | G | 2.6E-06 | 0.71 | 0.15 | 1.8E-07 | 0.71 | 0.14 | 0.1% | Distal |
| Intercellular Adhesion Molecule 1 (ICAM-1) | *ICAM1* | rs11575071 | 19 | 10272168 | G | C | 2.3E-45 | -0.49 | 0.04 | 3.4E-52 | -0.54 | 0.04 | 0.3% | Local |
|  | *ICAM1* | rs5491 | 19 | 10274864 | T | A | 2.5E-36 | -0.14 | 0.01 | 1.7E-17 | -0.10 | 0.01 | 4.5% | Local |
|  | *ICAM1* | rs139053442 | 19 | 10283720 | C | G | 9.1E-17 | -0.54 | 0.07 | 1.9E-17 | -0.53 | 0.06 | 0.1% | Local |
|  | *ICAM1* | rs28382777 | 19 | 10400963 | G | T | 6.6E-04 | -0.08 | 0.02 | 1.4E-09 | -0.13 | 0.02 | 0.6% | Local |
|  | *ICAM1* | rs5030400 | 19 | 10285120 | T | C | 4.7E-07 | 0.15 | 0.03 | 2.0E-07 | 0.14 | 0.03 | 0.4% | Local |
| Interleukin-6 (IL-6) | *IL6R* | rs568587329 | 1 | 154730517 | T | C | 5.4E-06 | -0.93 | 0.20 | 1.2E-06 | -0.99 | 0.20 | 0.0% | Distal |
| Lipoprotein-associated phospholipase A2 (Lp-PLA2) Activity | *PLA2G7* | rs144007943 | 6 | 46662909 | G | T | 8.0E-34 | -0.46 | 0.04 | 5.8E-36 | -0.48 | 0.04 | 0.2% | Local |
|  | *PLA2G7* | rs74479543 | 6 | 46784401 | A | G | 2.3E-22 | -0.13 | 0.01 | 1.8E-24 | -0.13 | 0.01 | 2.4% | Local |
|  | *PLA2G7* | rs144067869 | 6 | 46709433 | G | A | 1.7E-10 | -0.35 | 0.06 | 8.7E-11 | -0.36 | 0.06 | 0.1% | Local |
|  | *PLA2G7* | rs150641786 | 6 | 46774942 | A | C | 3.6E-03 | 0.05 | 0.02 | 1.1E-06 | 0.07 | 0.02 | 1.4% | Local |
|  | *APOE* | rs429358 | 19 | 44908684 | C | T | 4.2E-37 | 0.06 | 0.01 | 1.0E-13 | 0.06 | 0.01 | 13.7% | Distal |
|  | *APOE* | rs8106813 | 19 | 44928401 | G | A | 1.2E-02 | 0.01 | 0.00 | 1.2E-07 | 0.02 | 0.00 | 54.6% | Distal |
| Lipoprotein-associated phospholipase A2 (Lp-PLA2) Mass | *PLA2G7* | rs144007943 | 6 | 46662909 | G | T | 9.0E-25 | -0.39 | 0.04 | 5.1E-25 | -0.39 | 0.04 | 0.2% | Local |
|  | *PLA2G7* | rs74479543 | 6 | 46784401 | A | G | 2.6E-10 | -0.08 | 0.01 | 9.0E-13 | -0.09 | 0.01 | 2.4% | Local |
|  | *PLA2G7* | rs144067869 | 6 | 46709433 | G | A | 5.3E-11 | -0.39 | 0.06 | 2.5E-10 | -0.37 | 0.06 | 0.1% | Local |
|  | *PLA2G7* | rs73471140 | 6 | 46641939 | C | T | 3.1E-07 | -0.17 | 0.03 | 8.2E-09 | -0.19 | 0.03 | 0.3% | Local |
| P-selectin | *SELP* | rs6128 | 1 | 169593666 | T | C | 2.3E-10 | -0.05 | 0.01 | 5.8E-17 | -0.07 | 0.01 | 28.9% | Local |
|  | *SELP* | rs3917825 | 1 | 169595320 | G | A | 3.9E-07 | -0.19 | 0.04 | 2.9E-10 | -0.23 | 0.04 | 0.9% | Local |
|  | *SELP* | rs3917677 | 1 | 169622970 | C | A | 4.7E-09 | -0.31 | 0.05 | 5.1E-08 | -0.28 | 0.05 | 0.4% | Local |
|  | *ABO* | rs635634 | 9 | 133279427 | C | T | 1.0E-55 | 0.16 | 0.01 | 2.0E-15 | 0.19 | 0.02 | 84.7% | Distal |
| Matrix metalloproteinase-9 (MMP-9) | *MMP9* | rs3918249 | 20 | 46009497 | C | T | 1.6E-11 | 0.07 | 0.01 |  |  |  | 35.5% | Local |

Trait: trait name
Locus Name: significant loci identified by single variant analysis
rsID: rsID of lead signal
Unconditional: summary statistics of marginal analysis including p-value, beta coefficient, and standard error (SE)
EAF: Effect Allele Frequency (TOPMed), frequency of effect allele of the lead signal within TOPMed
Distal/Local: Distal means that the lead signal is more than 1Mb from the locus while local is in the 1Mb region on either side of the center of the locus

**Table 3.** Significant gene-centric and genetic region rare variant set analysis (after conditioning on known variants from the GWAS catalog and single-variant signals in the present analysis)

| Gene Centric Analysis |  |  |  |  |  |  |  |  |
| --- | --- | --- | --- | --- | --- | --- | --- | --- |
| Trait | CHR | Symbol | category | # variants | cMAC | STAAR-O p-value |  |  |
|  |  |  |  |  |  | unconditional | conditional | cond2round |
| C-Reactive Protein (CRP) | 1 | *CRP* | missense | 54 | 336 | 3.6E-22 | 1.3E-08 |  |
| Lipoprotein-associated phospholipase A2 (Lp-PLA2) Activity | 6 | *PLA2G7* | pLOF | 5 | 14 | 1.3E-13 | 1.1E-13 | 1.6E-06 |
|  | 6 | *PLA2G7* | missense | 56 | 323 | 6.4E-78 | 8.5E-28 | 3.9E-23 |
| Lipoprotein-associated phospholipase A2 (Lp-PLA2) Mass | 6 | *PLA2G7* | pLOF | 5 | 13 | 1.7E-10 | 1.1E-10 |  |
|  | 6 | *PLA2G7* | missense | 55 | 326 | 1.8E-75 | 1.1E-18 |  |

| Intercellular Adhesion Molecule 1 (ICAM-1) | 19 | *ICAM1* | missense | 69 | 451 | 7.8E-15 | 5.0E-08 | 3.9E-05 |
| --- | --- | --- | --- | --- | --- | --- | --- | --- |
|  | 19 | *ZNF653* | enhancer | 126 | 577 | 2.3E-11 | 2.3E-11 | 8.7E-01 |
| **Region-Based Analysis** |  |  |  |  |  |  |  |  |
| Trait | CHR | pos_min | pos_max | # variants | cMAC | STAAR-O p-value |  |  |
|  |  |  |  |  |  | unconditional | conditional | cond2round |
| Lipoprotein-associated phospholipase A2 (Lp-PLA2) Activity | 6 | 46707812 | 46709811 | 103 | 526 | 2.2E-46 | 1.7E-20 | 9.1E-10 |
|  | 6 | 46708812 | 46710811 | 95 | 389 | 6.9E-74 | 1.3E-21 | 4.2E-11 |
| Lipoprotein-associated phospholipase A2 (Lp-PLA2) Mass | 6 | 46707812 | 46709811 | 103 | 532 | 5.3E-44 | 9.3E-14 |  |
|  | 6 | 46708812 | 46710811 | 94 | 394 | 1.3E-64 | 2.0E-14 |  |
| Intercellular Adhesion Molecule 1 (ICAM-1) | 19 | 11282547 | 11284546 | 68 | 591 | 6.1E-12 | 4.5E-10 | 6.8E-01 |
|  | 19 | 11283547 | 11285546 | 91 | 892 | 7.7E-12 | 5.6E-10 | 9.7E-01 |
|  | 19 | 11284547 | 11286546 | 96 | 1337 | 1.8E-09 | 7.0E-09 | 6.5E-01 |
|  | 19 | 11285547 | 11287546 | 85 | 871 | 1.1E-09 | 1.8E-08 | 7.4E-01 |
|  | 19 | 11503547 | 11505546 | 119 | 729 | 2.8E-11 | 2.8E-11 | 8.3E-01 |

Trait: trait name
CHR: chromosome where the gene is located
Symbol: gene symbols
Category: category of gene-based test; pLOF means putative loss of function
pos\_min: starting position of the region tested, hg38
pos\_max: ending position of the region tested, hg38

**Table 4.**
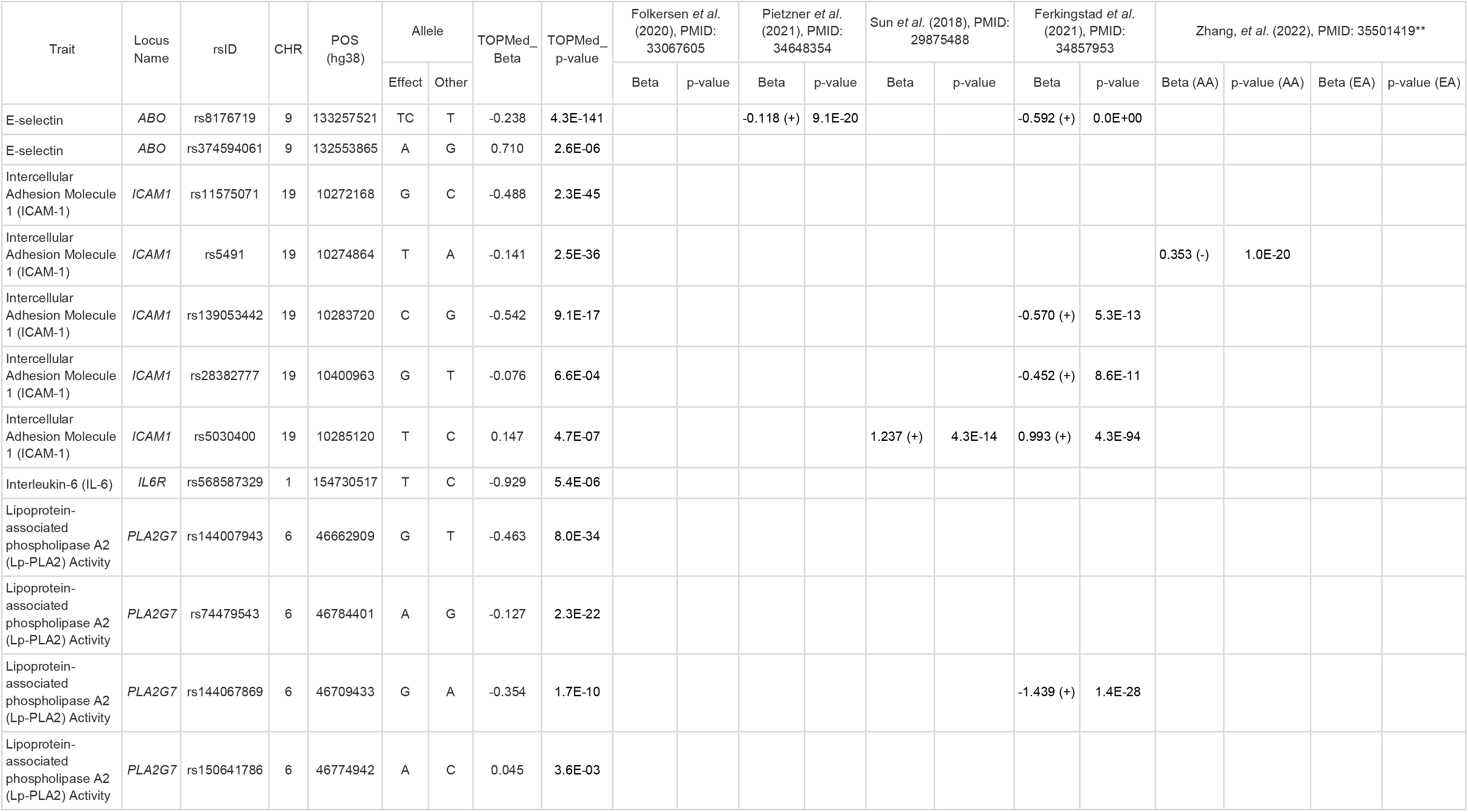
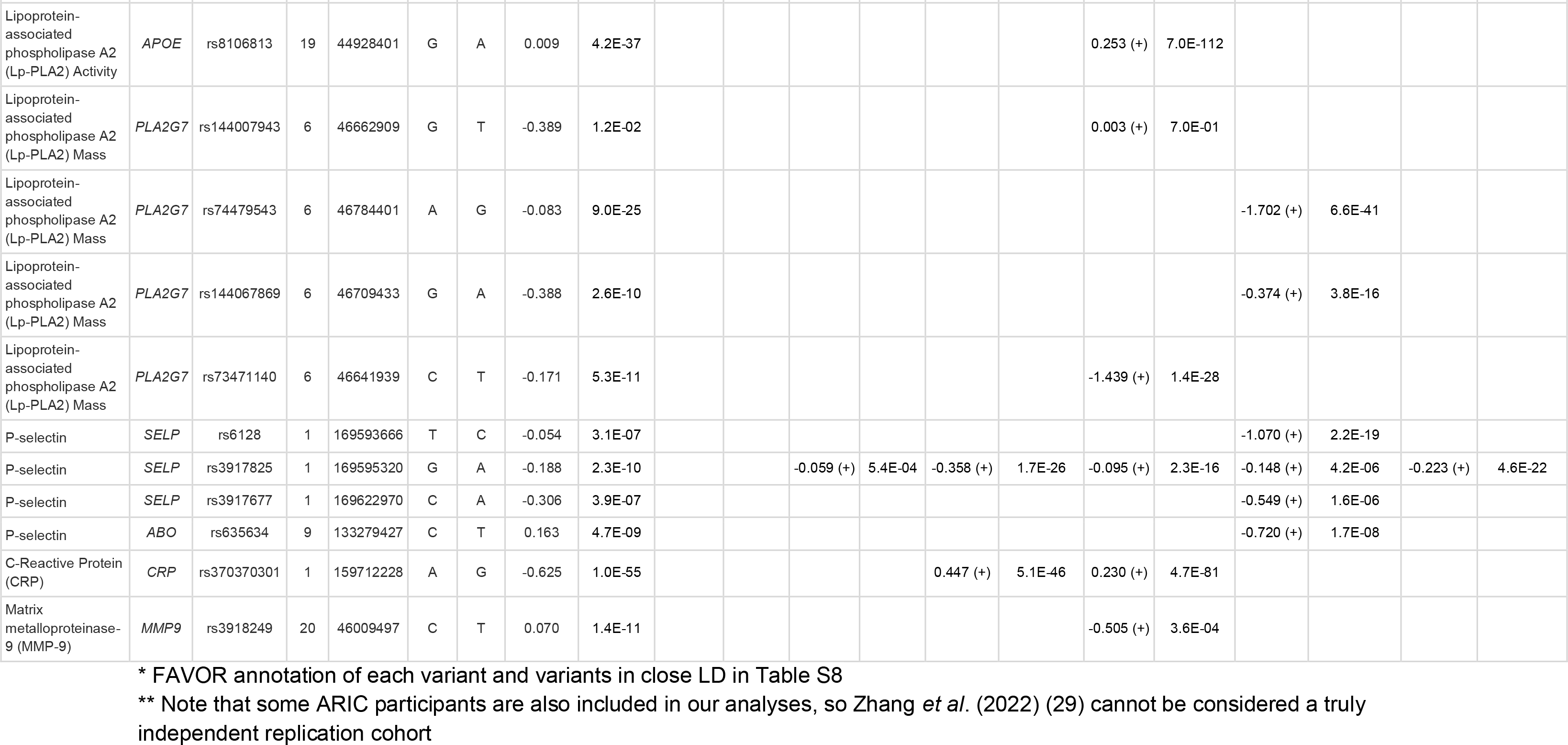
Replication of newly identified signals in previous semiquantitative platform pQTL analysis*.

### C-Reactive Protein

We identified genetic variants associated with CRP consistent with and expanding upon our previous analysis of CRP in 23,279 TOPMed participants (17). All 8 distinct single variant signals at the *CRP* locus previously known in TOPMed (17) (in partially overlapping samples) were also found here, and we identified 1 additional distinct signal, rs370370301. This non-coding, rare variant (TOPMed Effect Allele Frequency (EAF): 0.2%, 1000G EUR EAF: 0.1%, 1000G SAS EAF: 0.1%, and not available in all other populations in 1000G) did not reach genome-wide significance in the previous TOPMed analysis (p=5.0×10^-6^) but was associated with p=1.4×10^-11^ in the present analysis, with the smaller p-value likely attributable to the increased sample of 38,465 participants. This variant was not previously identified in an analysis of CRP in UK Biobank (UKB) (22), likely due to the rare frequency, but was identified in protein quantitative trait loci (pQTL) analysis of CRP measured using an untargeted proteomics platform by Ferkingstad *et al*. (2021) (Table 4, see Replication section in Methods below). Rare variant analysis yielded 1 significantly associated gene-centric set of 54 missense rare variants (p=3.6×10^-22^) on *CRP* locus driven in part by rs77832441 (p=7.8×10^-16^ for analysis of individual variant in TOPMed) (Tables S10B). We also tested a similar gene-centric missense rare variant set for association in UKB (p=6.4×10^-34^ based on 116 variants, details in Table S15). rs77832441 (MAC=153, EAF=0.2%) was previously identified in Schick *et al*. (25), and other subthreshold CRP associated missense variants have been identified in the CARDIA study (26).

We note that rs77832441 was pruned from the conditional analysis list based on LD (see Methods) but a variant in close LD, rs553202904 (r^2^=0.97), was included (Tables S6, S10C and S11C), and the significance of the gene centric test was attenuated but still at least nominally significant (missense set, p=1.3×10^-8^, Tables 3 and S10A) when this signal was adjusted for, suggesting additional subthreshold CRP missense variants in particular remain to be identified as individually significant in larger analyses.

In addition to signals at the *CRP* locus, we also identified multiple loci in the single variant association analyses not previously detected in prior TOPMed analysis, including 3 with multiple distinct signals (*LEPR*, *SALL1*, *APOE*) (Table S5). Each of these signals were attenuated below the genome-wide significance threshold after adjusting for known associations from the GWAS catalog and other prior publications (17,22). We also attempted to replicate single variant findings from semi-quantitative proteomics platforms (Table S13). Many of the previously reported pQTL lead signals were also found by our marginal single variant association analysis (with p<0.05 and the same direction of effect), including 10 out of 11 available CRP lead signals from Ferkingstad *et al*. (2021) (26), 5 out of 5 CRP lead signals from Pietzner *et al*. (2021) (27), 2 out of 2 available CRP lead signals from Sun *et al*. (2018) (28), 4 out of 5 CRP lead signals from African American Atherosclerosis Risk in Communities (ARIC) participants, and 5 out of 5 CRP lead signals from European American ARIC participants from Zhang *et al*. (2022) (29). Note that some ARIC participants are also included in our analyses, so this is not an independent replication sample for Zhang *et al*. (2022) (29) findings. Similar look-ups were performed for all other overlapping traits and are noted in Table S13.

### E-selectin

There are 9 distinct signals at the *SELL/SELE*, *FUT6*, and *ABO* loci associated with E-selectin, and 2 distinct signals remaining at the *ABO* locus after conditioning on previously identified signals, including single variant signals from previous TOPMed analysis. This pair of signals, rs8176719 and rs374594061, were the second and third distinct signals in our marginal analysis. Variant rs8176719 is a frameshift insertion exonic variant common across all populations that tags blood group O (30). We do note that in our prior work from TOPMed (18), while this was not captured as an independent genome-wide signal, associations of differential E-selectin levels across blood groups (with O treated as reference) were also observed. This variant’s association with E-selectin further illustrates the extensive pleiotropy of the *ABO* locus, which has been previously associated with diseases such as malaria, venous thromboembolism, and COVID-19 and traits such as vWF and Factor VIII levels. This association was also identified by Ferkingstad *et al*. (2021) (26) and Pietzner *et al*. (2021) (27) using high throughput semiquantitative proteomics platforms (Table 4) (27). E-selectin associated distinct variant rs374594061 is rare across all populations (TOPMed EAF: 0.9%, and not available in 1000G) and, likely as a consequence, has no previously reported associations in the GWAS catalog and was also not tested in available replication cohorts.

### Intercellular adhesion molecule 1

For ICAM-1, we identified 9 distinct single variant signals at the *ICAM1* and *ABO* loci; 5 distinct signals at *ICAM1* remained after conditioning on known associations (Table S7). The GWAS conditionally significant association at rs5491, the fourth distinct signal in unconditional results at the *ICAM1* locus, is an exonic variant (TOPMed EAF: 4.5%, 1000G AFR EAF: 25.0%, 1000G AMR EAF: 1.7%, 1000G EAS EAF: 5.3%, 1000G EUR EAF: 0.7%, 1000G SAS EAF: 2.0%) that is low frequency in most populations but common among African ancestry populations. We do note that prior work has found assay-binding artifacts for coding variants in *ICAM1* (31); this variant and its LD proxies were not *ICAM1* eQTLs in eQTLGen phase I (32) and Genotype-Tissue Expression (GTEx) V8 (33) look-ups (as described in Methods) and we suspect it may be an assay interference effect. There are 4 other conditionally distinct noncoding variants – rs11575071, rs139053442, rs28382777, rs5030400 – at the ICAM1 locus (Table 2); most are low frequency or rare across all populations. rs5030400 was identified as a distinct signal in both Sun *et al*. (2018) (28) and Ferkingstad *et al*. (2021) (26). rs11575071, rs139053442, and rs28382777 were identified as distinct signals in Ferkingstad *et al*. (2021) (26). As displayed in Figure S17, there is some long-range LD for variants identified in the *ICAM1* locus, notably for rs5491 (displayed in turquoise) in Figure S17.

We also identified multiple conditionally significant rare variant set-based associations with ICAM-1 including 2 gene-centric sets (Table S10A) and 6 2-kb sliding windows (Table S11A, individual variants included in tests included in Table S11B), and 2 of them overlap the *ICAM1* locus. We identify a set of missense rare variants at *ICAM1*, whose most significant variant was the identified rs139053442 association but which remains significant after conditioning on rs139053442 and other single variant findings from TOPMed and other studies (Table S6); it includes additional variants such as rs5030400 which was also identified by Sun *et al*. (2018) (28) and Ferkingstad *et al*. (2021) (26) using semiquantitative proteomics data from SomaScan for ICAM-1.

### Matrix Metalloproteinase-9

We identified the *MMP9* encoding gene for association with MMP-9 levels in single variant analysis. This cis pQTL locus included 1 distinct signal at intronic variant rs3918249 that was common in all populations, and it has repressed regulatory function with high H3K27me3 score 48 according to FAVOR (20). This variant was also identified by Ferkingstad *et al*. (2021) (26), Pietzner *et al*. (2021) (27), and Sun *et al*. (2018) (28), but to our knowledge this is the first report using a quantitative immunoassay. Our identified variant rs3918249 (Figure 1, TOPMed EAF 35.5%) is highly linked (r^2^=0.938) with coding variant rs17576 (Table S8). Similar to rs5491 for ICAM-1 and other coding variants, it is possible such a coding variant signal may tag an antibody binding effect without true impact on protein abundance. However, we note that rs3918249 is also highly linked with rs6017721 (r^2^=0.86) and rs4810482 (r^2^=0.92), both of which are significant conditionally distinct lead variants in GTEx V8 cis-eQTL results for MMP-9 (Table S9). The finding suggests that this variant influences transcript and likely protein abundance, not just antibody binding to the MMP-9 target protein. The MMP-9 coding variant rs3918249 we identified is in moderately LD (r^2^=0.664) with the intronic variant rs3918253. rs3918253 is associated with liver enzyme levels; this close linkage disequilibrium suggests MMP-9 abundance could mediate this liver-related signal rs3918253 (34). Notably, our identified variant is not in LD (r^2^=0.037) with MMP-9 coding variant rs2250889, which was identified in analysis of MMP-9 levels on SOMAscan in the INTERVAL study (28) and recent proteomic analyses in Icelandic populations (26). rs2250889 is nominally associated with MMP-9 in TOPMed (p=5.5×10^-3^) in marginal analysis, but not significant (p=0.16) after conditioning on rs3918249.

### P-selectin

For P-selectin, we identified 5 distinct single variant signals at the *SELP* locus (Table S5), and 3 of them remain significant after conditioning on known associations (Table S7), and 1 distinct single variant signal at the *ABO* locus that is significant conditional on known associations (Tables S5 and S7). At the *SELP* locus, 2 of 3 conditionally significant signals are intronic (rs3917677, rs3917825). rs3917825 is relatively conserved (top 9.1% genome-wide aPC-conservation score) (20). Both of these variants are low frequency in AFR ancestry participants (1.7% for rs3917677 in 1000G, 2.8% for rs3917825 in 1000G) and not observed in EUR ancestry participants (from reference panels). The remaining significant signal in the *SELP* locus is the synonymous variant rs6128, which is more common in AFR ancestry (53.3%) than in EUR ancestry (16.6%) participants from 1000G. Variant rs6128 is a platelet splice QTL that alters SELP exon 14 skipping and soluble versus transmembrane P-selectin protein production (35). Although rs6128 was not reported in the GWAS catalog for ELISA-measured P-selectin, it was previously identified in the INTERVAL study using the SOMAscan assay platform (28), and also identified by Ferkingstad *et al*. (2021) (26), and Pietzner *et al*. (2021) (27).

For aggregate tests of rare variants, lead signals were detected at 2 consecutive 2-kb sliding windows in the *SELP* locus located at chr1:169615464-169617463 and chr1:169616464-169618463 (Table S11A), which are driven in part by rs7529463. This coding variant is highly conserved (top 1.6% genome-wide aPC-conservation score), very rare (TOPMed AF 0.1%), and has high aPC protein function scores (top 0.2% genome wide) (20).

At the *ABO* locus, the distinct signal (rs635634, which tags blood group A) remained significant after conditioning on known variants (Table S6); however, the p-value is significantly attenuated (from p=1.0×10^-55^ to p=2.0×10^-15^, Table S7) when adjusting for known GWAS catalog variants. This variant was also identified by Ferkingstad *et al*. (2021) (26), Pietzner *et al*. (2021) (27), and Sun *et al*. (2018) (28). As might be anticipated given the widespread pleiotropy of the *ABO* locus, rs635634 is also related to many other traits in the GWAS catalog, including cholesterol (36–39), CRP (40), type 2 diabetes (41), and blood cell phenotypes such as white blood cell count (42).

### Interleukin 6

We identified 1 distinct signal for IL-6 at the *IL6R* locus in single variant analysis, variant rs61812598 (p=1.1×10^-49^, Table S5). After conditioning on previous GWAS-identified variants (Table S6), this initial distinct signal was no longer significant (p=4.9×10^-6^); however, a new signal at rs568587329 was identified (Table S7). This variant was subthreshold in both the marginal analysis (p=5.4×10^-6^, Table S5) and conditional analysis (p=4.7×10^-6^, Table S5) adjusting for sentinel variant rs61812598 of the *IL6R* locus. The association of rs568587329 was modestly strengthened when adjusted for known variants at the *IL6R* locus (GWAS catalog, (43)) (p=1.2×10^-6^, Table S7). This variant is rare in all populations (TOPMed EAF: 0.03%, 1000G AFR EAF: 0.4%, and not available in all other populations in 1000G) and has a high aPC-Transcription-Factor score 17.29 (top 1.87% genome wide) (20).

### Lipoprotein-Associated Phospholipase A2 activity and mass

For the Lp-PLA2 activity trait, we identified 11 distinct single variant signals at the *CELSR2, APOE, LDLR,* and *PLA2G7* loci (Table S5). After conditioning on previous GWAS identified variants (Table S6), 2 GWAS conditional distinct signals remain at the *APOE* locus (Table S7), and 4 GWAS conditional distinct signals remain at the *PLA2G7* locus (Table S7).

At the *APOE* locus, the GWAS conditional distinct signals rs429358 (representing the well-known APOE-ε4 allele) and rs8106813 are the second and the third distinct signals of our stepwise analysis. rs429358 is common across all populations and was also identified by Ferkingstad *et al*. (2021) as associated with Lp-PLA2(26). Reflective of known pleiotropy at the *APOE* locus, rs429358 has been associated with many other traits, including Alzheimer’s disease (44–57), cholesterol (39,58–64), red cell distribution width (65), liver enzyme levels (34), blood protein levels (13,28), and CRP levels (5,40,59,63,64,66,67), including in the present analysis. rs8106813 was also reported to be related to Alzheimer’s disease (68).

We observe 4 distinct signals, rs144007943, rs74479543, rs144067869, and rs150641786, at the *PLA2G7* locus significant upon conditioning on prior GWAS identified signals. Each of these variants are rare, and only rs144067869 was identified in prior semi-quantitative proteomics efforts (26). In addition to these single variant associations, we observe 2 gene centric and 2 2-kb sliding windows significantly associated at the *PLA2G7* locus. We observe a set of putative loss-of-function (pLOF) rare variants and missense rare variants. The pLOF set is partly driven by rs140020965, whereas the missense set is partly driven by rs200303358 (though the set is still quite significant after conditioning on this variant (p=3.9×10^-23^) (Tables S10A). We also observe a 2-kb sliding window spanning chr6:46707812-46709811 and another 2-kb sliding window spanning chr6:46708812-46710811 both partially driven by rs140020965 and rs200303358 (Tables S11A).

For Lp-PLA2 mass, we identified 6 distinct signals at the *PLA2G7* locus (Table S5). After conditioning on previous GWAS-identified variants (Table S6), 4 signals remained significant (Table S7) – rs144007943, rs74479543, rs144067869, and rs73471140 – 3 of which were identified in our analysis of Lp-PLA2 activity, unsurprisingly given the high correlation between the traits. The additional signal at rs73471140 is rare across all populations and in very low LD with all Lp-PLA2 activity lead variants (r^2^<0.01). We again observe associations with pLOF rare variants and missense rare variants at the *PLA2G7* locus (Table S10A), and the same 2 significant 2-kb sliding windows as Lp-PLA2 activity (spanning chr6:46707812-46709811 and chr6:46708812-46710811) are also significant.

## Discussion

We sought to evaluate the genetic determinants of 21 inflammation biomarkers using data from the TOPMed Program. Previous efforts in TOPMed with E-selectin (18) and CRP (17) demonstrated that inclusion of diverse cohorts yielded further insights into the genetic determinants of these biomarkers. Our work extends these findings by incorporating both larger samples for these previously analyzed traits and expanding the scope to include 19 additional traits and rare variant aggregate tests. We identified significant associations with 6 traits in single variant analysis and 5 traits in aggregate rare variant analysis that remained significant after conditioning on known associations.

Our findings demonstrate the complementary value of performing both single and rare variant analyses when analyzing quantitative traits. Recent analyses of quantitative lipid traits from TOPMed also combined single and rare variant analyses, similarly finding both common signals and conditionally distinct aggregate rare variant signals, mostly at known genes, for both coding and noncoding variant sets (69), similar to our findings here. Several exome sequencing efforts for diverse traits and diseases, for example waist hip ratio (70) and schizophrenia (71), have similarly identified joint impacts from common noncoding variants and rare coding variants at the same loci (including at Mendelian genes), but similar findings in the noncoding space have been less widely reported. Previous analysis (17) of CRP in TOPMed identified variants in enhancer regions (including 1 whose impact on transcription and protein binding to the enhancer region was validated *in vitro*) that were more common in AFR versus EUR ancestry individuals, demonstrating the contributions of ancestry differentiated variants in noncoding regions to the genetic architecture of the trait. That analysis did not include aggregate tests of rare variants, and in the present analysis we observe that even after conditioning on known single variant associations additional signals are identified by performing aggregate analyses. We identify a similar joint contribution of common, rare, and low frequency variants for multiple traits, including P-selectin and ICAM-1. We do note that in some cases our rare variant signals are consecutive or overlapping, suggesting that multiple rare signals within a broad region may contribute to gene regulation (Lp-PLA2 and ICAM-1). We note that it remains an outstanding challenge to completely disentangle whether a common or rare variant signal is driving biological processes, and continued large-scale analysis will likely provide further insight.

Our analysis yielded more distinct signals than previously detected for inflammation biomarkers, primarily at known loci. This finding points to the extensive allelic heterogeneity at, in particular, encoding gene loci, as reflected by the increased number of statistically distinct cis pQTL (26) and cis eQTL (33) distinct signals observed with increasing sample size. Studies of populations with different ancestry often observe different cis eQTL and pQTL signals due to ancestry differentiated allele frequencies for such variants (72,73), including our own analyses of CRP within TOPMed (17). Prior work suggested that such distinct signals can have different molecular mechanisms (even acting through distinct transcripts, as at the adiponectin encoding gene locus (74)), with variants in different distinct signals often impacting different regulatory regions (including distinct enhancer and promoter regions). We anticipate that expanded efforts to understand such “secondary” distinct signals at known GWAS identified loci for quantitative traits in expanded sample sizes will identify many additional loci with significant allelic heterogeneity and ancestry differentiated QTLs. Such analyses would be completed ideally with individual level data to avoid issues with approximate conditional analysis with poor matching between the LD reference panel and the GWAS or WGS analysis population. Both individual level sequence data and improved imputation reference panels (23,75,76) may help increase discovery in the low frequency/rare variant space.

Our analysis further highlights the value of including study populations inclusive of multiple ancestry groups. Using a larger sample size, we confirmed findings from previous TOPMed analyses driven by variants common only in AFR reference populations including rs3917422 and rs17855739 for E-selectin (18), as well as rs11265259 and rs181704186 for CRP (17). Given the diversity of our sample, we were able to additionally identify associations with Lp-PLA2 traits, P-selectin, and ICAM1 that were exclusively or disproportionately observed in AFR reference populations (Fig 1). Many previous large-scale analyses have been conducted primarily in European ancestry individuals.

Coding cis pQTLs present particular challenges for biomarker traits. Such QTLs often have large effect sizes, but it is unclear whether these effects represent a true impact on protein abundance versus interference with antibody/aptamer binding. Such issues have also been identified in previous work from TOPMed, notably for the E-selectin signal rs3917422 identified by Polfus *et al*. (2019) (18), as well as in prior genetic analyses for other antibody measured biomarker traits (31,77,78). As a supplemental analysis, we assessed coincidence of our identified coding pQTL signals with distinct eQTL signals in GTEx V8 (33) and eQTLGen phase I (32), and found that our MMP-9 coding variant signal, but not the signal at ICAM-1, coincided with an eQTL. When such coding pQTL variants also influence transcription, it is less likely they are an aptamer or antibody effect. This should be carefully evaluated in future pQTL efforts, using both quantitative and semiquantitative platforms.

There are multiple limitations of our present analysis. While the TOPMed program provides a rich sequencing data source, there are a relatively limited set of cohorts within TOPMed that have measured inflammation biomarkers in their participants. Similarly, few other large scale studies have incorporated inflammation biomarker measurement, and most of those have primarily limited their measurements to CRP (22). This limits our ability to perform a well-powered analysis among some traits in TOPMed, and to replicate our findings in external datasets. To partially address this limitation, we conducted single variant replication analyses using semi quantitative proteomics platforms, and in general note good replication rates for our single variant findings – for variants tested in both datasets, 16 out of 18 variants are both significant and in the same direction between previous semi-quantitative pQTL analysis and our TOPMed analysis (Table 4). We also replicate many distinct signals from prior semiquantitative high throughput platform publications in our immunoassay-based findings – for variants tested in both datasets, 217 out of 431 variants are both significant and in the same direction between previous semi-quantitative pQTL analyses and our TOPMed analysis (Table S13). Correlation both between ELISAs themselves and between ELISA and aptamer assays (as well as between Olink and SomaScan) varies, and will impact expected replication rates (79–81). However, such information is unfortunately not available for the vast majority of the specific immunoassays used here. We also note that many of our biomarkers are still mostly measured in non-Hispanic White participants; future efforts should focus on further increasing the inclusion of additional populations.

Through our analysis of 21 inflammation biomarkers, we identified additional signals and highlighted features of such large-scale analyses. Across this set of traits, consistently observed features included a combination of contributing common and rare variant signal, extensive allelic heterogeneity, and ancestry specificity of some identified variants. Such features have been observed in other efforts, such as the analysis of lipids and blood cell traits in the TOPMed program (82). We anticipate that with continually increasing sample sizes (and thereby statistical power) that these key aspects of our study would be observed in similar sequencing-based analyses of complex traits.

## Methods

### Whole Genome Sequencing

We analyzed variants with whole genome sequencing from blood in samples from the NHLBI TOPMed program. All participants had deep coverage sequencing, with harmonization, variant discovery, and genotype calling previously described (82). We specifically leveraged data from Freeze 8, which was aligned to GRCh38 reads (83). All positions in this manuscript are reported based on GRCh38. Samples were processed by the TOPMed Data Coordinating Center, resulting in 1.02B variants for 138K samples. For all Freeze 8 samples, population principal components of genetic ancestry were calculated using PC-AiR (84), genetic relatedness was calculated using PC-Relate (85), and race/ethnicity was reported by each study (mostly from participant self-report). Full single variant and aggregate test summary statistics will be provided at time of publication to the TOPMed genomic summary result dbGaP accession (phs001974).

### Phenotype Harmonization and Study Sample

Phenotype harmonization for 21 inflammation biomarkers was primarily performed by the TOPMed Data Coordinating Center (86) as previously described. COPDGene, GeneSTAR, and WHI were harmonized directly from study-provided data. Methods of inflammation biomarker measurement are listed in Table S1. We note that not all cohorts used the same platform, and samples run on multiple platforms are not available for assay re-calibration. This is unfortunately a common limitation for cross-cohort analyses of inflammation biomarker traits. Study participants were included based on informed consent restrictions (excluding some individuals with consent for only disease specific analyses), duplicates were removed to retain observations with the highest frequency assay type where applicable, trait measurements exceeding 3 standard deviations from the mean were removed, and individuals with missing data were excluded. CRP was natural log-transformed to address non-normality in distribution. All traits were analyzed after rank-based inverse normal transformation, performed by study-race/ethnicity strata, with variance rescaled within each strata. The present analysis of inflammation biomarkers included sample sizes ranging from 737-38,465 individuals from 12 cohorts in Freeze 8 of the NHLBI TOPMed program. Across all traits, the sample is primarily non-Hispanic White, though efforts were made to include a multi-ethnic population wherever possible. The sample is described in Tables S1 and S3.

### Single Variant Analysis

We performed single variant analyses across ancestry groups as was done in several previous studies in TOPMed (17,87–90). We tested PASS variants (based on support vector machine variant classifier, as previously described in TOPMed sequencing methods (82) with a minor allele count (MAC) of at least 10 in our pooled sample, resulting in a test of between 11,793,614 - 57,072,499 variants for each biomarker trait. We used linear mixed effects models (91) as implemented in GENetic Estimation and Inference in Structured samples (GENESIS 2.19.1 (92)) on the BioData Catalyst Seven Bridges platform (93), adjusting for age, sex, variables combining study and race and ethnicity, an empirical kinship matrix for relatedness and population structure, 11 ancestry principal components (84,85) and permitting heterogeneous variance across the strata of the combination of study and race and ethnicity. Differences in ancestry were accounted for by our principal components and kinship matrix adjustment, and we also adjusted for race/ethnicity as a self or study reported variable, given previously reported impacts of these social constructs on levels of inflammatory biomarkers (94,95). Loci were defined as statistically significant according to a genome-wide threshold given as 1×10^-9^ (21).

We next performed stepwise conditional analysis at significant loci to identify the total number of conditionally distinct signals within a +/- 1 Mb (+/- 3Mb for *ICAM1* chr19) window. Conditional analysis was performed by running the association analysis conditioning on the lead variant defined by p-value, and repeating this process until no variants were significant at the locus. Significance was defined at alpha=0.05 using a Bonferroni correction for the number of variants tested within the locus, for example 0.05/39,488=1.3×10^-6^ for CRP at the *CRP* locus. The threshold for conditional analysis of each trait conditioning on distinct signals and known variants are listed in Tables S5 and S7.

### Identification of Distinct Signals Through Conditional Analysis

Many previous studies of inflammatory biomarkers have identified genome-wide significant signals for the inflammation biomarkers tested here (Table S6). To identify which single variant signals in our analysis were distinct from previously identified GWAS variants, we performed stepwise conditional analysis at significant loci for each trait, conditioning on the reported associations from the GWAS Catalog, Raffield *et al*. (2020) (17), Sinnott-Armstrong *et al*. (2021) (22), Ahluwalia *et al*. (2021) (43), Folkersen *et al*. (2017) (96), and Polfus *et al*. (2019) (18) as covariates in our null model to determine which associations in our TOPMed analysis are distinct from those previously identified. We mapped published associations within a +/- 1Mb window (+/- 3Mb window for ICAM1 chr19 due to very long range LD) of the TOPMed identified loci (i.e. GWAS conditional distinct signals at Table S7) to TOPMed Freeze 8 variants by positions and alleles. To avoid collinearity, we pruned the previous GWAS identified variant set with the linkage disequilibrium threshold r^2^=0.9 to obtain a list of previously identified distinct signals at each locus. All known variants were included as fixed effects in the null model. If any variants were still significant using a locus-wide threshold after this adjustment for known variants, we proceeded to perform stepwise conditional analysis again, to identify the total number of distinct signals after adjustment for known variants from prior GWAS.

### Rare Variant Analysis

We performed rare variant analysis for both gene-centric and genetic region aggregation units. We tested PASS variants with MAC at least 1 and minor allele frequency (MAF) less than 1.0% in our pooled sample. We used linear mixed effects models with weighting by functional annotation as implemented in STAAR (97–99), adjusting for age, sex, race/ethnicity-study, and 11 population ancestry principal components and permitting heterogeneous variance across race-study strata and empirical kinship for relatedness and population structure. Gene-centric units were defined for all protein-coding genes using coding annotations based on GENCODE consequences as (a) putative loss of function (stop gain, stop loss, splicing), (b) missense, and (c) synonymous variants; non-coding variants were captured via masks characterized by (a) promoters if within +/- 3kb of a transcription start site overlayed with DHS signal, or (b) enhancers if identified by GeneHancer overlayed with DHS signal. Genetic region analysis used 2-kb sliding windows with a 1kb skip length.

The STAAR-O p-value, incorporating 2 weighting schemes using the beta distribution based on MAF (with α_1_ = 1, α_2_ = 25 to upweight rarer variants or with α_1_ = α_2_ = 1 treat all equally) in addition to annotation-based weights using CADD, LINSIGHT, FATHMM-XF, aPC-Protein-Function, aPC-Conservation, aPC-Epigenetics-Active, aPC-Epigenetics-Repressed, aPC-Epigenetics-Transcription, aPC-Local-Nucleotide-Diversity, aPC-Mutation-Density, aPC-Transcription-Factor, aPC-Mappability, aPC-Proximity-To-TSS-TES, was considered. Sets were defined as statistically significant according to a Bonferroni-corrected significance threshold separately for gene-centric, correcting for all 5 masks, and genetic region analysis, correcting for all windows (Table S12). We performed conditional analysis to identify signals by obtaining trait-specific associations from the GWAS catalog and the single-variant analysis in a 1 Mb window from the start and end of the positions spanned by the set.

### Annotation

We used multiple resources to obtain functional annotations for inclusion in the rare variant analysis and to describe identified variants, including FAVOR, GTEx, and ANNOVAR. We obtain aPCs from FAVOR (20,97), providing summarized functional categories by aggregating correlated individual functional annotations. These aPCs provide variant-level measures as a PHRED score yielding the interpretation that scores greater than 10 within a given functional category are in the top 10% for all TOPMed variants.

### Replication

Many genetic loci and distinct signals have been identified in previous pQTL studies using untargeted semiquantitative platforms (SomaScan and Olink) (26–29,100). For our conditionally distinct signals (GWAS conditional distinct signals at known loci, and rs3918249 for MMP-9), we pulled results from summary statistics of these prior published studies and compared their direction of effect and level of significance with our findings in TOPMed (Table 4). Conversely, we also attempted to replicate all previously reported distinct pQTL signals for overlapping traits in our summary statistics (Table S13).

For the CRP phenotype, we replicated our results using 188,912 samples with whole genome sequencing data from UKB (23,24). The null model was constructed using the same methods as the TOPMed analyses, and both single variant and variant set analyses were conducted using STAARPipeline app (https://github.com/xihaoli/staarpipeline-rap) (97,98) on the UKB Research Analysis Platform (RAP).

### eQTL Coincidence

We also checked the coincidence of eQTL signals from cis-eQTLGen phase I (32) and GTEx V8 (33) for the distinct signals we detected on the corresponding coding region of the inflammation biomarker traits. For cis-eQTLGen, we performed GCTA-COJO (101) on the summary-based Mendelian randomization (102) formatted cis-eQTLGen results to identify statistically distinct lead signals. For GTEx V8 (33), conditionally distinct signals were already reported (details in Table S9).

## Supporting information

Supplemental Figures and Acknowledgements

## Acknowledgements

Molecular data for the TOPMed program was supported by the National Heart, Lung and Blood Institute (NHLBI). Study-specific omics support information can be found in the supplement. Core support including centralized genomic read mapping and genotype calling, along with variant quality metrics and filtering, were provided by the TOPMed Informatics Research Center (3R01HL-117626-02S1; contract HHSN268201800002I). Core support including phenotype harmonization, data management, sample-identity QC, and general program coordination, were provided by the TOPMed Data Coordinating Center (R01HL-120393; U01HL-120393; contract HHSN268201800001I). We gratefully acknowledge the studies and participants who provided biological samples and data for TOPMed. TOPMed specific acknowledgments for studies are included in Table S2. Additional study specific acknowledgments are included under cohort descriptions in the Table S1.

Support for this work was provided by the National Institutes of Health, National Heart, Lung, and Blood Institute, through the BioData Catalyst program (awards 1OT3HL142479-01, 1OT3HL142478-01, 1OT3HL142481-01, 1OT3HL142480-01, and 1OT3HL147154).

The authors wish to acknowledge the contributions of the consortium working on the development of the NHLBI BioData Catalyst® (BDC) ecosystem. LMR, SG, and ZL were supported by NHLBI BioData Catalyst Fellowship program. XLI was supported by NHLBI TOPMed Fellowship program. LMR was also supported by R01HG010297. XLIN was supported by NHLBI R01HL163560 and NHGRI U01 HG009088 and U01HG012064. The project described was also supported by the National Center for Advancing Translational Sciences, National Institutes of Health, through Grant KL2TR002490 (LMR). The content is solely the responsibility of the authors and does not necessarily represent the official views of the NIH.

The Genotype-Tissue Expression (GTEx) Project was supported by the Common Fund of the Office of the Director of the National Institutes of Health, and by NCI, NHGRI, NHLBI, NIDA, NIMH, and NINDS. The data used for the analyses described in this manuscript were obtained from the GTEx Portal on 03/31/2020.

## Notes

### Competing Interest Statement

SMG is now affiliated with Regeneron Genetics Center but her affliation is correct when she performed analysis for this work.
LMR is a consultant for the TOPMed Administrative Coordinating Center (through Westat).
Psaty serves on the Steering Committee of the Yale Open Data Access Project funded by Johnson & Johnson.
In the past three years, Edwin K. Silverman received grant support from Bayer and Northpond Laboratories.
XLin is a consultant of AbbVie Pharmaceuticals and Verily Life Sciences.

