## Supplemental Figures and Acknowledgements for "Whole Genome Sequencing Based Analysis of Inflammation Biomarkers in the Trans-Omics for Precision Medicine (TOPMed) Consortium"

### Table of Content

Fig S1. Overview of 21 inflammation-related biomarkers.
Fig S2. Lead signals of CRP on *CRP*.
Fig S3. Lead signals of CRP on *LEPR*.
Fig S4. Lead signal of CRP on *IL6R*.
Fig S5. Lead signal of CRP on *NLRP3*.
Fig S6. Lead signal of CRP on *GCKR*.
Fig S7. Lead signal of CRP on *ILF10*.
Fig S8. Lead signal of CRP on *IL6*.
Fig S9. Lead signal of CRP on *PPP1R3B*.
Fig S10. Lead signal of CRP on *HNF1A*.
Fig S11. Lead signals of CRP on *SALL1*.
Fig S12. Lead signals of CRP on *APOE*.
Fig S13. Lead signal of E-selectin on *SELE*.
Fig S14. Lead signal of E-selectin on *ABO*, prior to and after conditioning on previous literature reported variants.
Fig S15. Lead signal of E-selectin on *FUT6*.
Figure S16. Lead signal of ICAM-1 on *ABO*.
Figure S17. Lead signals of ICAM-1 in *ICAM1*, prior to and after conditioning on previous literature reported variants.
Fig S18. Lead signals of MMP-9 on *MMP9*.
Figure S19. Lead signals of P-selectin on *SELP*, prior to and after conditioning on previous literature reported variants.
Fig S20. Lead signal of P-selectin on *ABO*, prior to and after conditioning on previous literature reported variants.
Fig S21. Lead signals of IL-6 on *IL6R*, prior to and after conditioning on previous literature reported variants.
Fig S22. Lead signals of Lp-PLA2 Activity on *CELSR2*.
Fig S23. Lead signals of Lp-PLA2 Activity on *PLA2G7*, prior to and after conditioning on previous literature reported variants.
Fig S24. Lead signals of Lp-PLA2 Activity on *APOE*, prior to and after conditioning on previous literature reported variants.
Fig S25. Lead signals of Lp-PLA2 Activity on *LDLR*.
Fig S26. Lead signals of Lp-PLA2 Mass on *PLA2G7*, prior to and after conditioning on previous literature reported variants.
Fig S27. Lead signals of IL-18 on *NLRC4*.
Fig S28. Lead signals of IL-18 on *IL18*.
Fig S29. Lead signal of MCP-1 on *ACKR1*.
Fig S30. Lead signal of MCP-1 on *CCR2*.
Fig S31. Lead signal of TNFR2 on *TNFR2*.
Fig S32. QQ plots of marginal single variant association analysis of each trait.
Cohort Acknowledgements

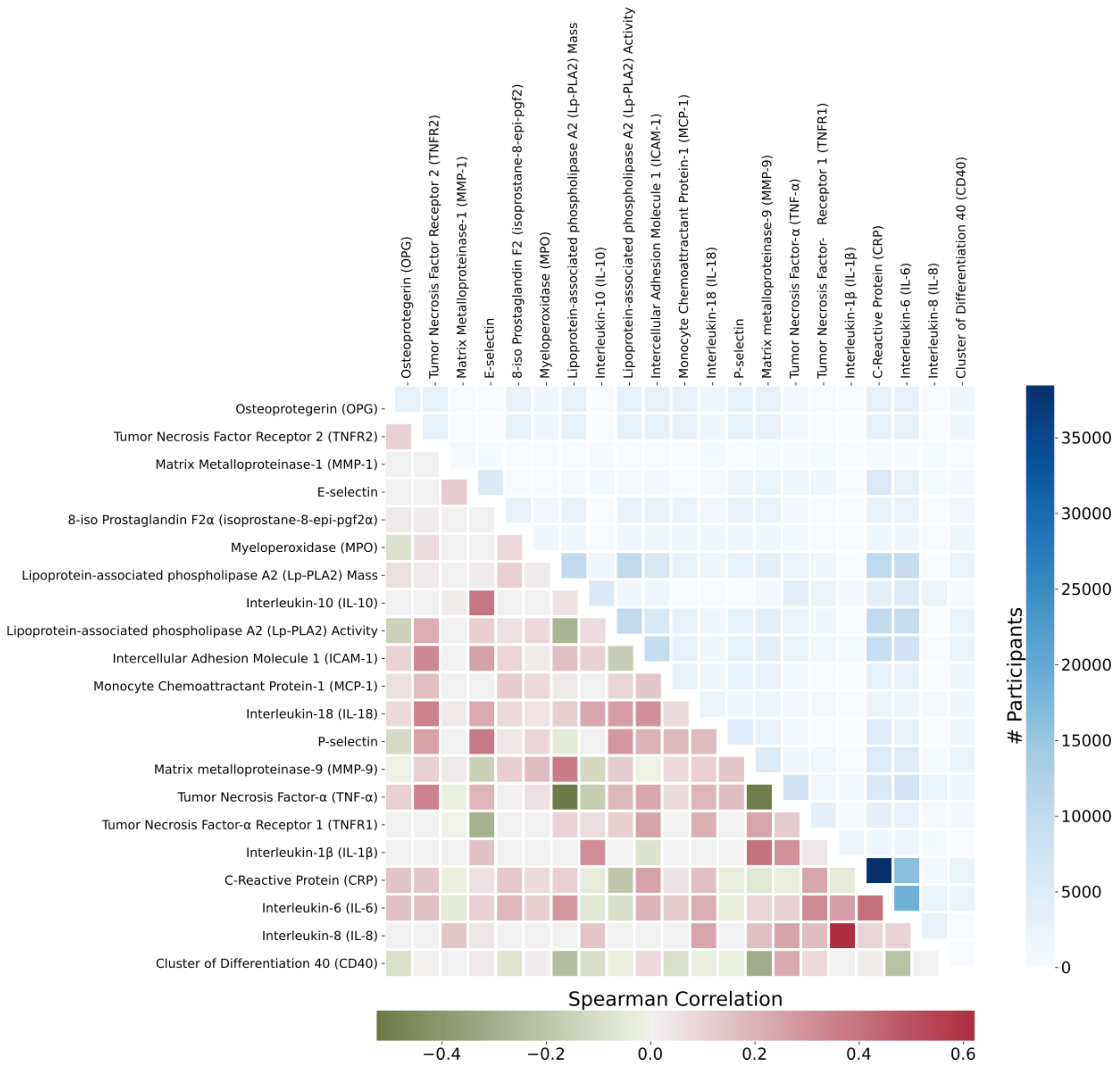


**Fig S1. Overview of 21 inflammation-related biomarkers.** The upper triangle panel: (1) the diagonal cells show the number of participants of each trait; (2) the off-diagonal cells show the number of overlapped participants for any pair of traits. Details are in Table S14. The lower triangle panel is the correlation heatmap of 21 biomarkers. Correlation is measured by Spearman correlation between the rank-based inverse normalized protein abundance of the shared individuals in any pair of traits.The most correlated pair of traits are IL-8 and IL-1β (r=0.622) – IL-8 was measured in 2,826 individuals, and IL-1β was measured in 1,638 individuals, while the number of overlapped individuals is 334. The most negative correlated pair of traits are TNF-α and Lp-PLA2 mass (r=-0.524) – TNF-α was measured in 7,591 individuals, and Lp-PLA2 mass was measured in 10,132 individuals, while the number of overlapped individuals is 2,419. The correlations of other pairs of traits are less than 0.408 and larger than -0.509.

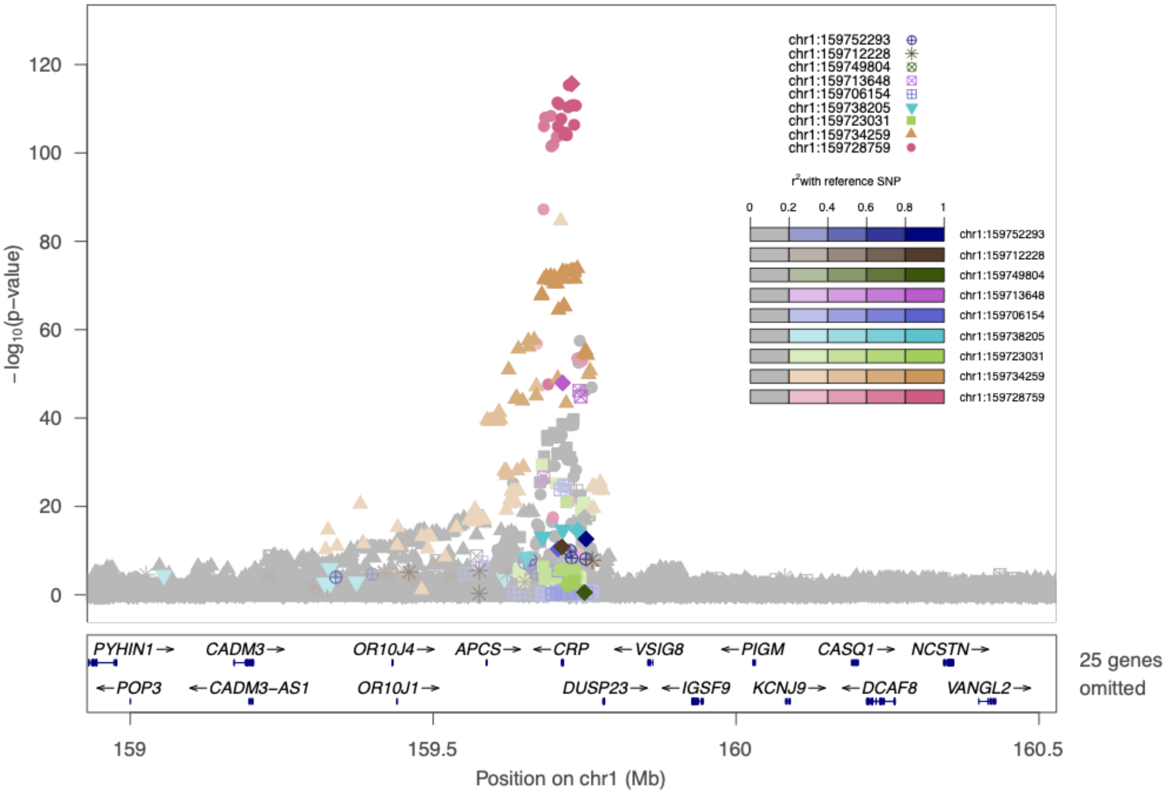


**Fig S2. Lead signals of CRP on *CRP*.** Lead signals are chr1:159728759_A_G (rs7553007), chr1:159734259_A_G (rs59330569), chr1:159723031_G_A (rs2794518), chr1:159738205_G_A (rs553202904), chr1:159706154_C_T (rs11265259), chr1:159713648_G_C (rs1800947), chr1:159749804_T_A (rs12734907), chr1:159712228_A_G (rs370370301), and chr1:159752293_G_A (rs181704186).

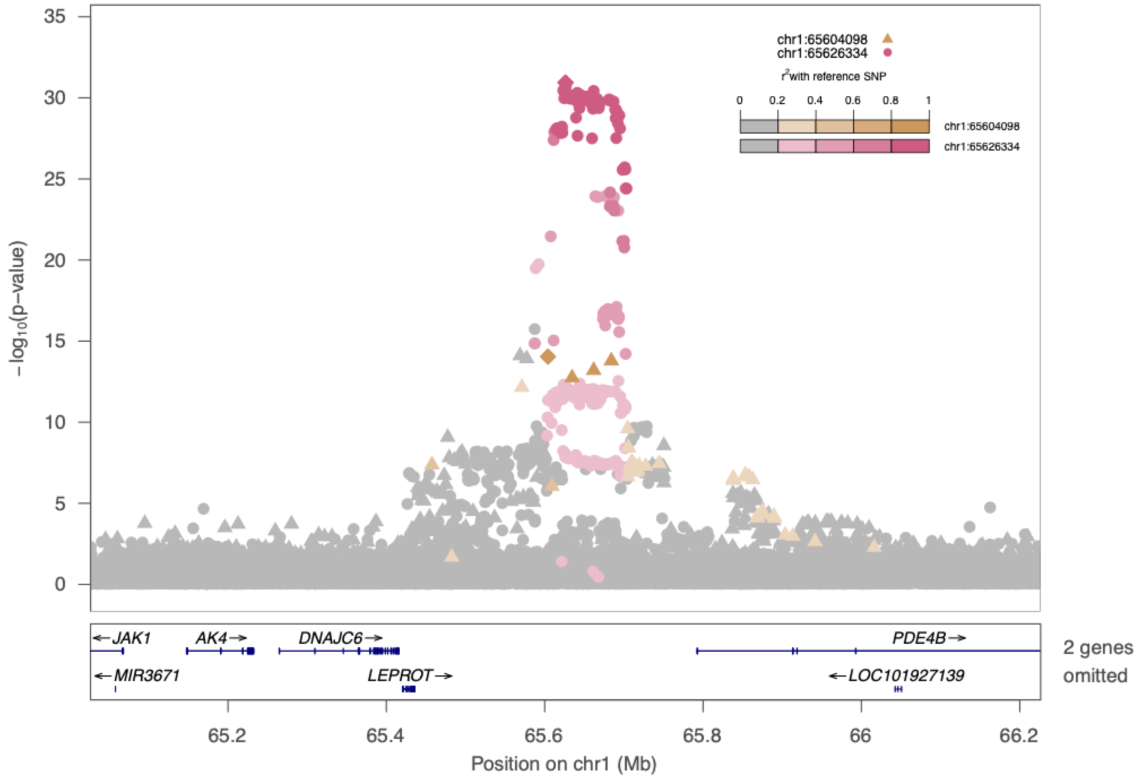


**Fig S3. Lead signals of CRP on *LEPR*.** Lead signals are chr1:65626334_A_T (rs6588153) and chr1:65604098_G_A (rs72683129).

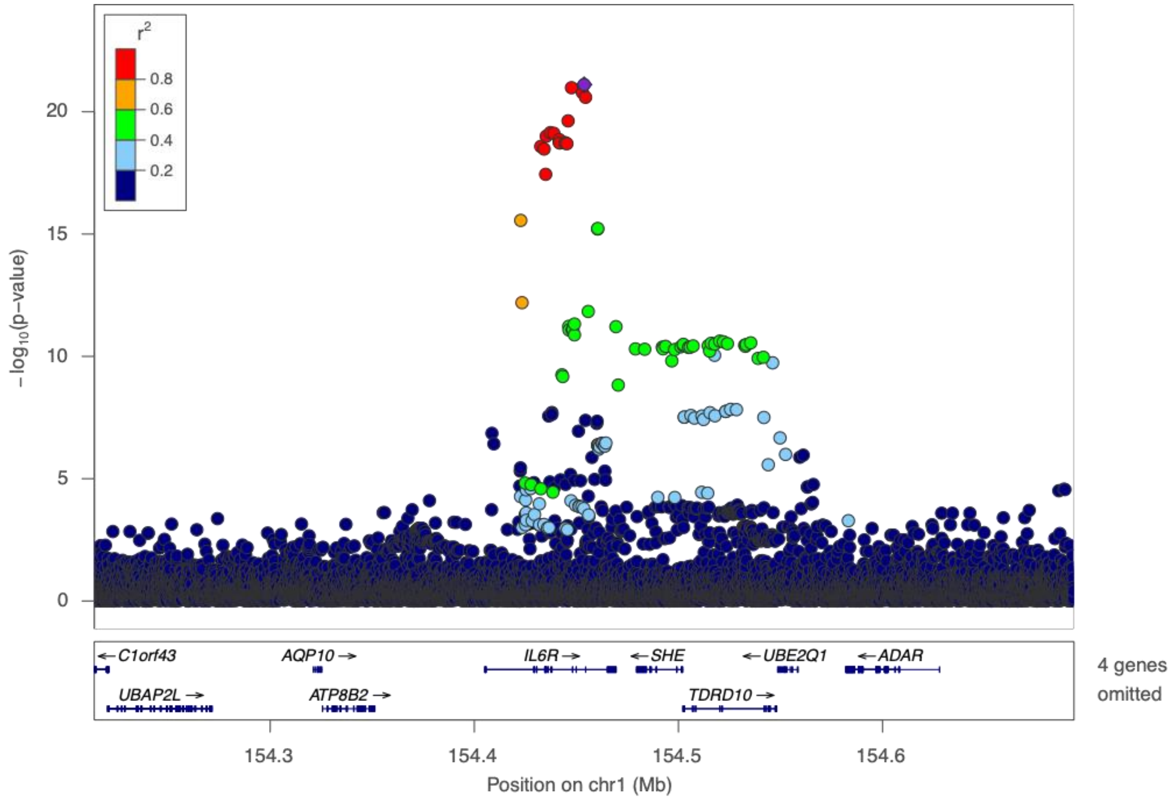

**Fig S4. Lead signal of CRP on *IL6R*.** Lead signal is chr1:154453788_C_T (rs4129267).

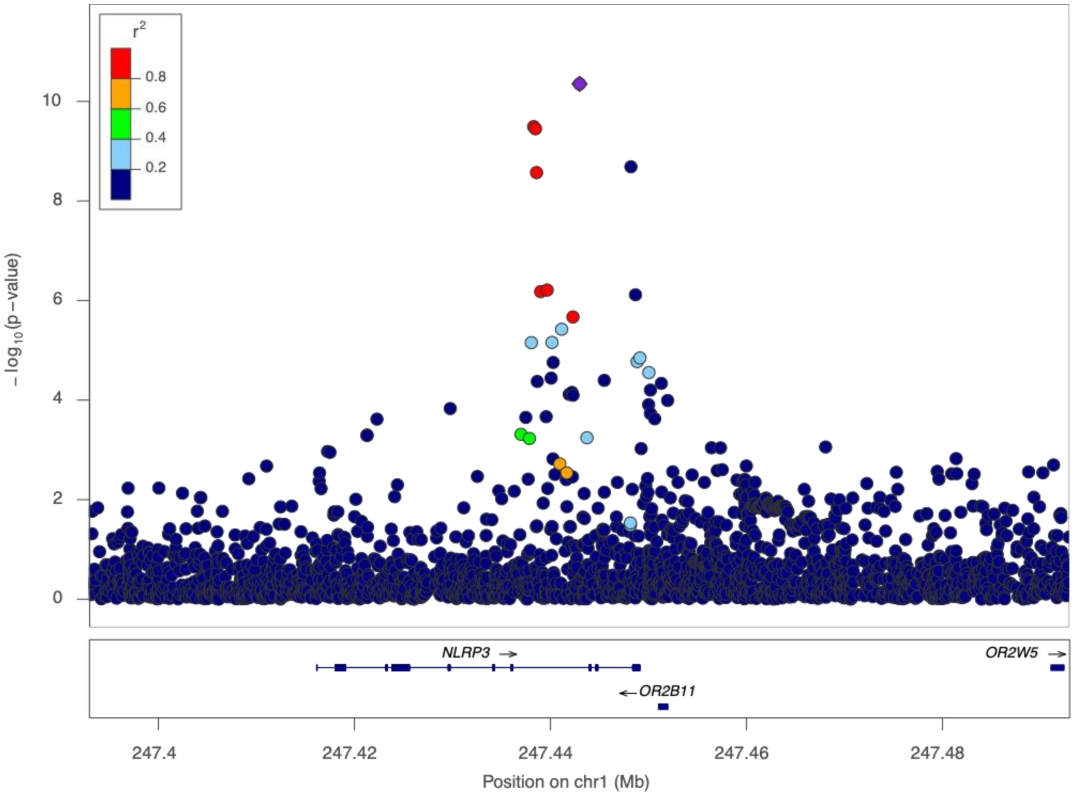


**Fig S5. Lead signal of CRP on *NLRP3*.** Lead signal is chr1:247442974_T_C (rs56188865).

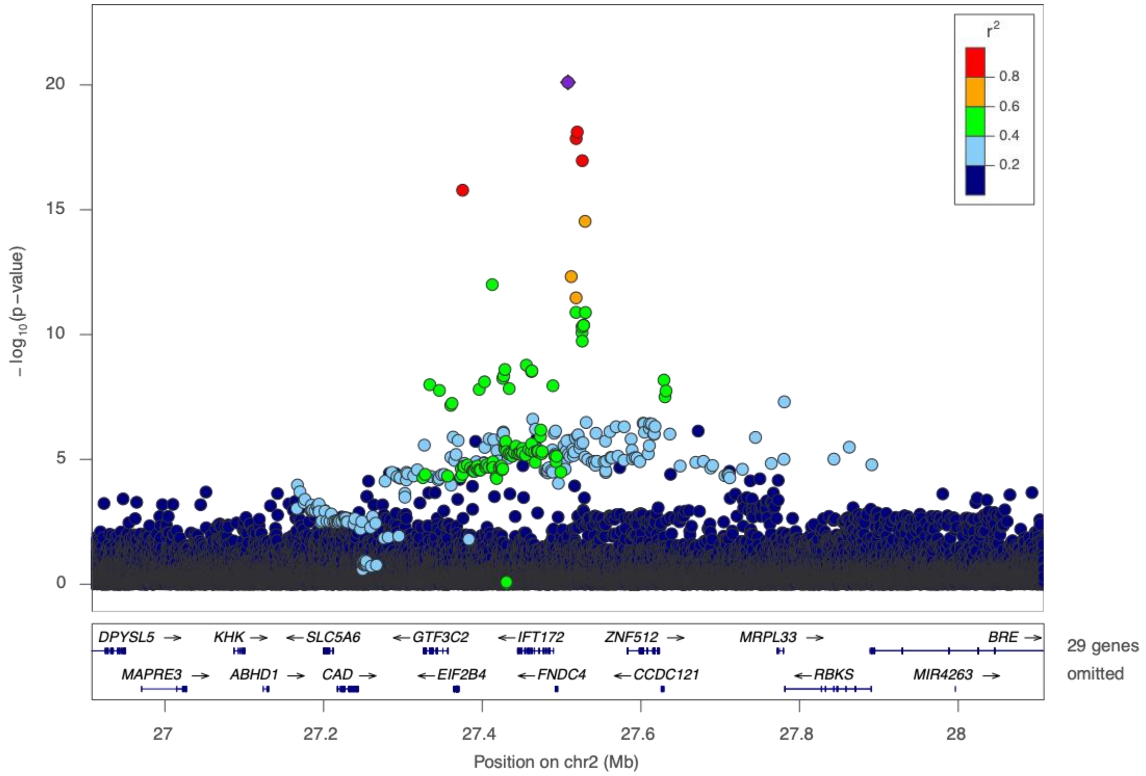


**Fig S6. Lead signal of CRP on *GCKR*.** Lead signal is chr2:27508073_T_C (rs1260326).

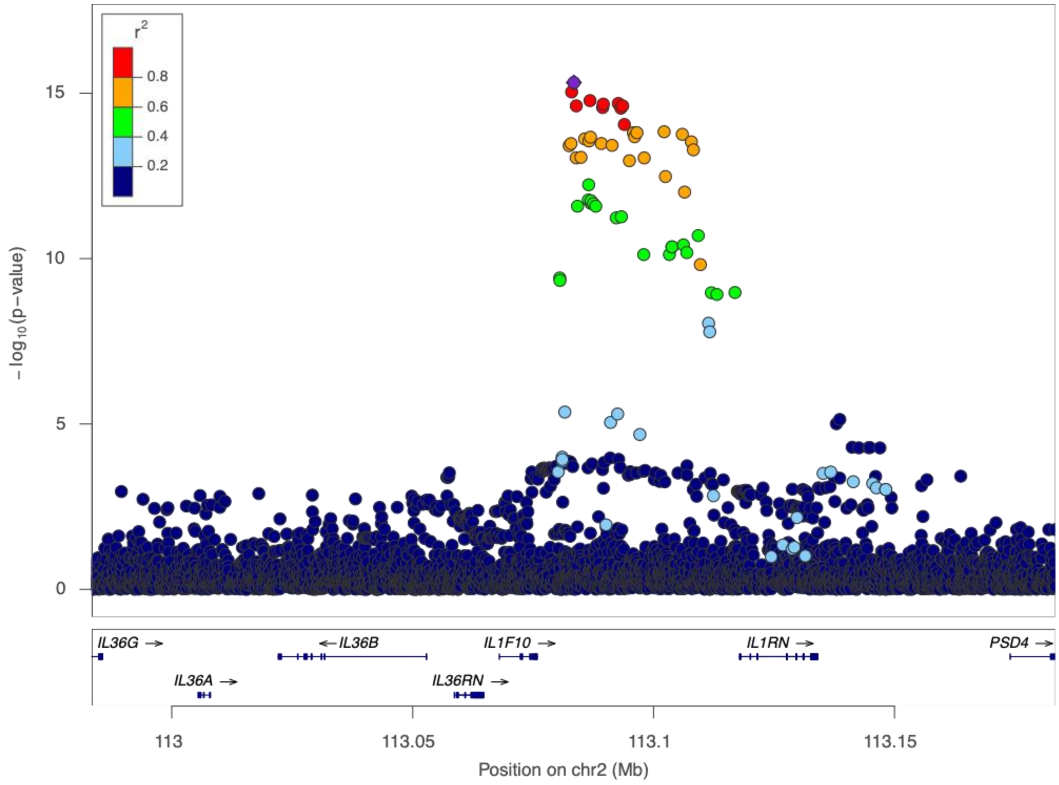

**Fig S7. Lead signal of CRP on *ILF10*.** Lead signal is chr2:113083453_A_G (rs6734238).

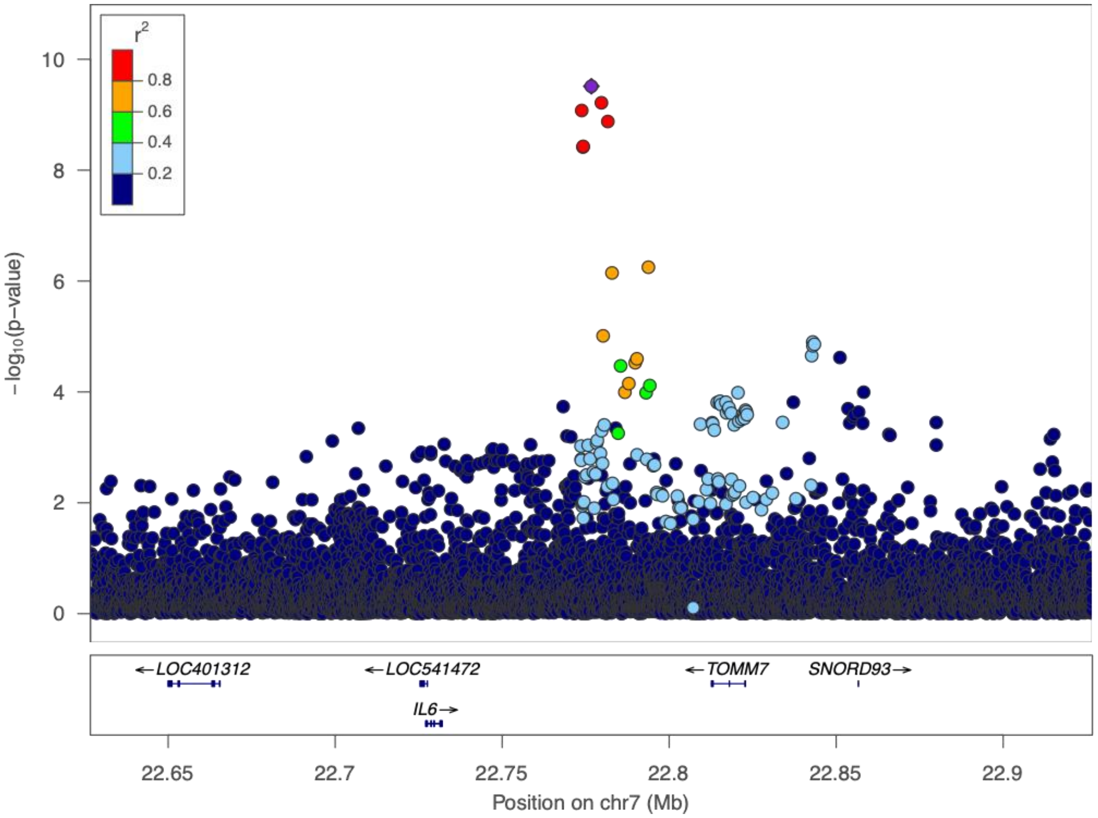


**Fig S8. Lead signal of CRP on *IL6*.** Lead signal is chr7:22776697_A_G (rs10279140).

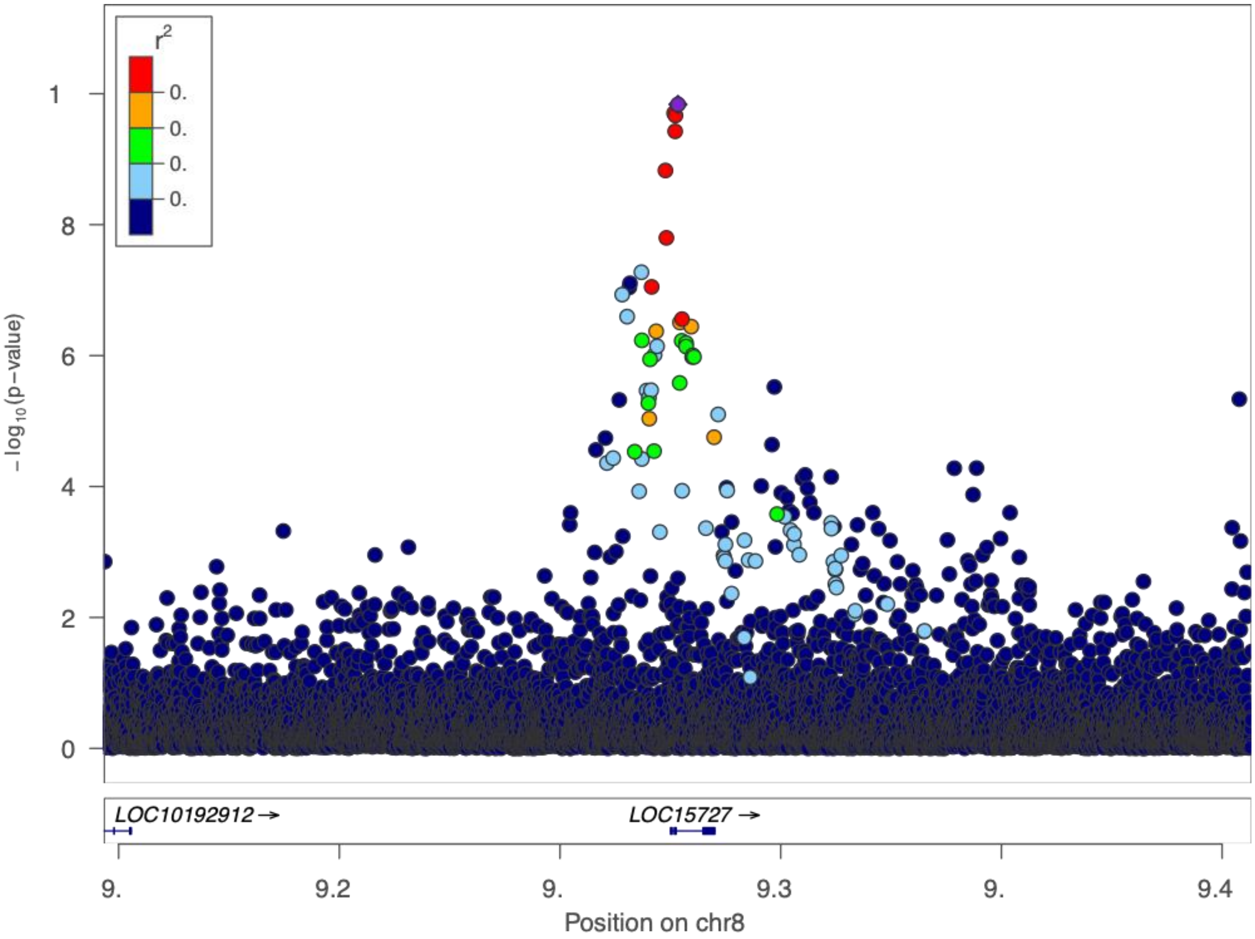


**Fig S9. Lead signal of CRP on *PPP1R3B*.** Lead signal is chr8:9326721_G_A (rs4240624).

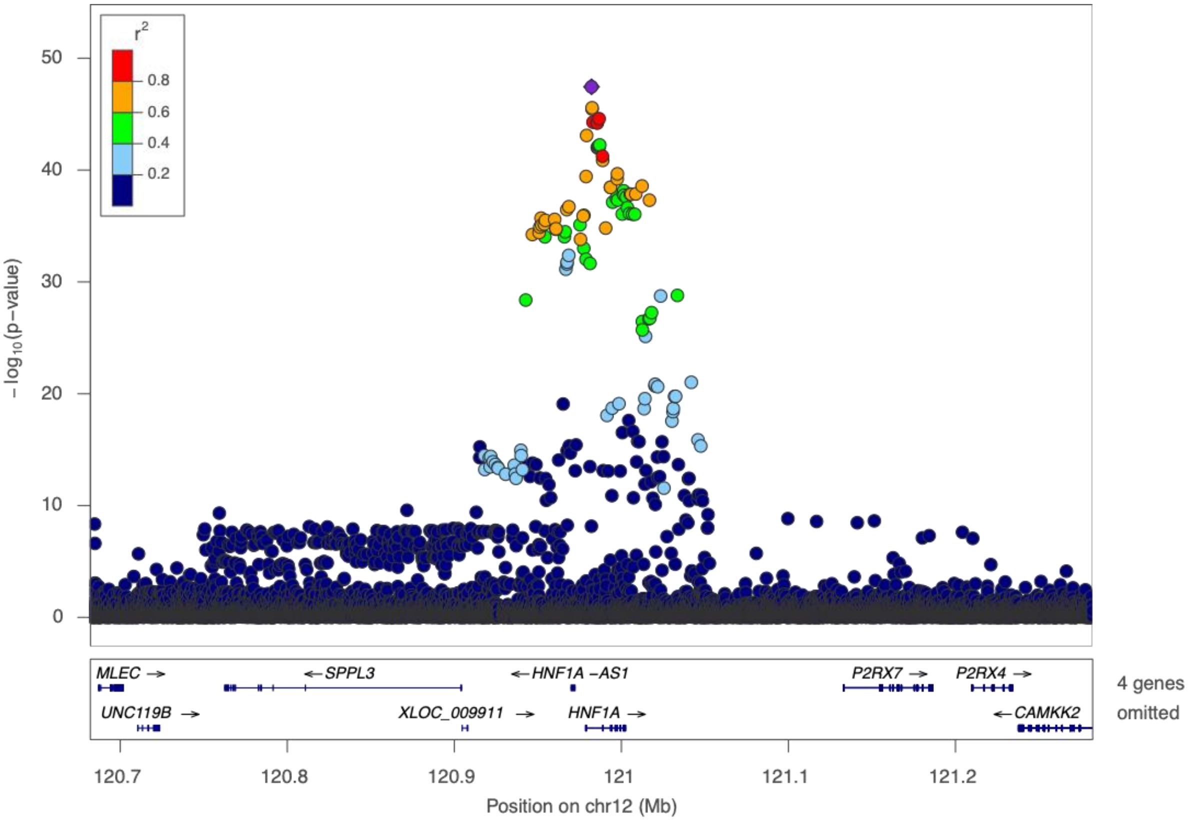


**Fig S10. Lead signal of CRP on *HNF1A*.** Lead signal is chr12:120982123_T_C (rs1169284).

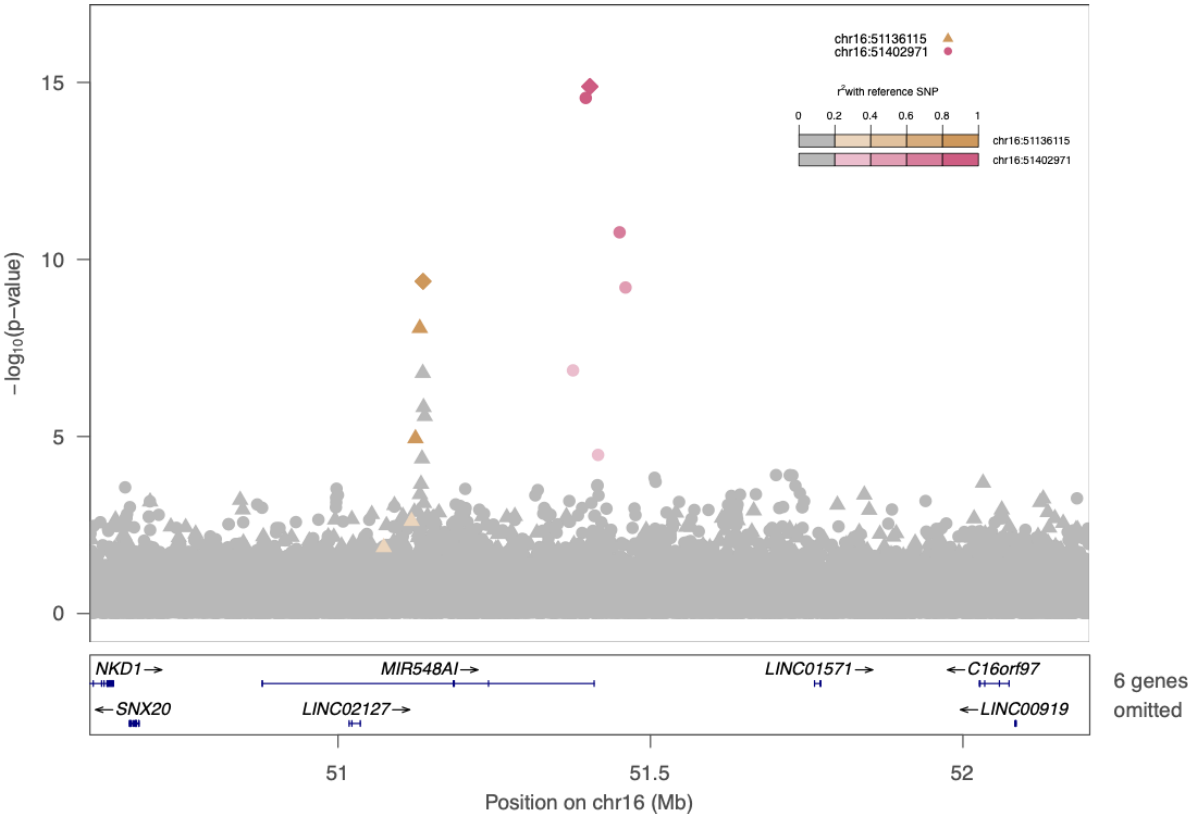


**Fig S11. Lead signals of CRP on *SALL1*.** Lead signals are chr16:51402971_A_G (rs17616063) and chr16:51136115_G_T (rs116971887).

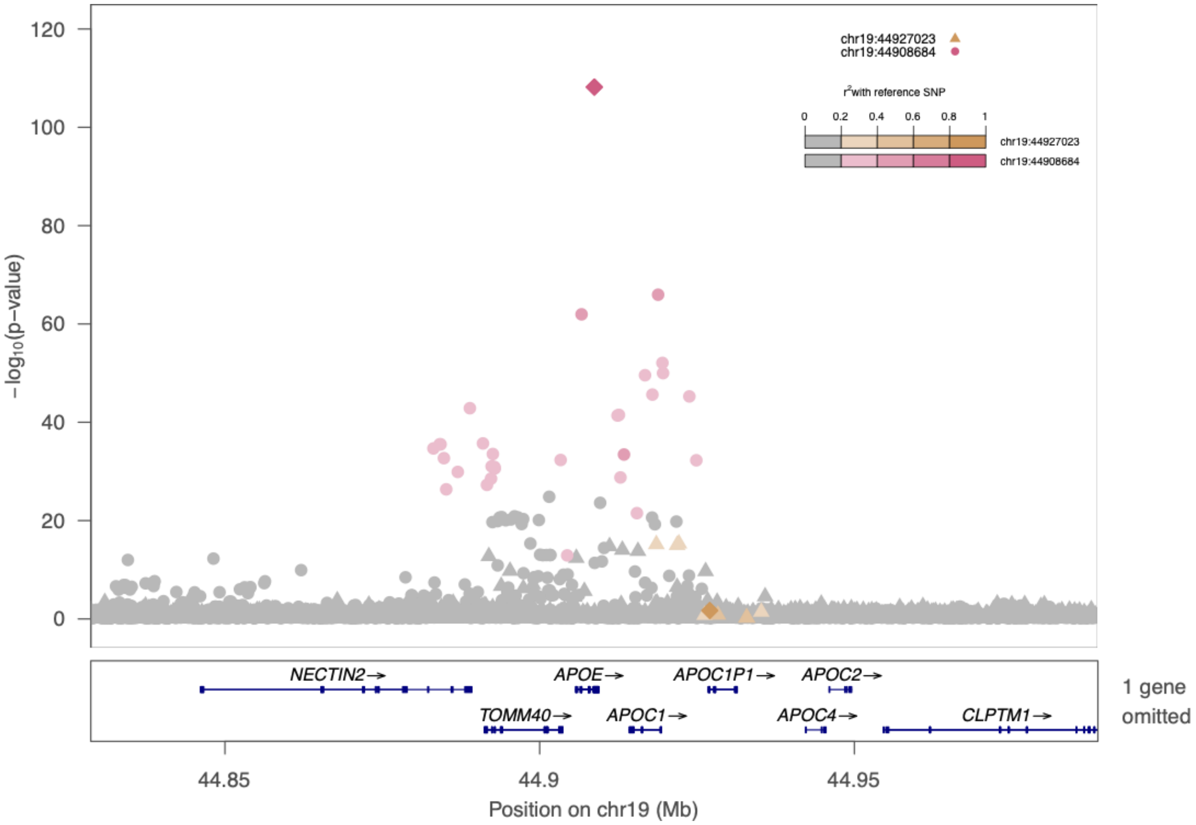


**Fig S12. Lead signals of CRP on *APOE*.** Lead signals are chr19:44908684_T_C (rs429358) and chr19:44927023_C_G (rs5112).

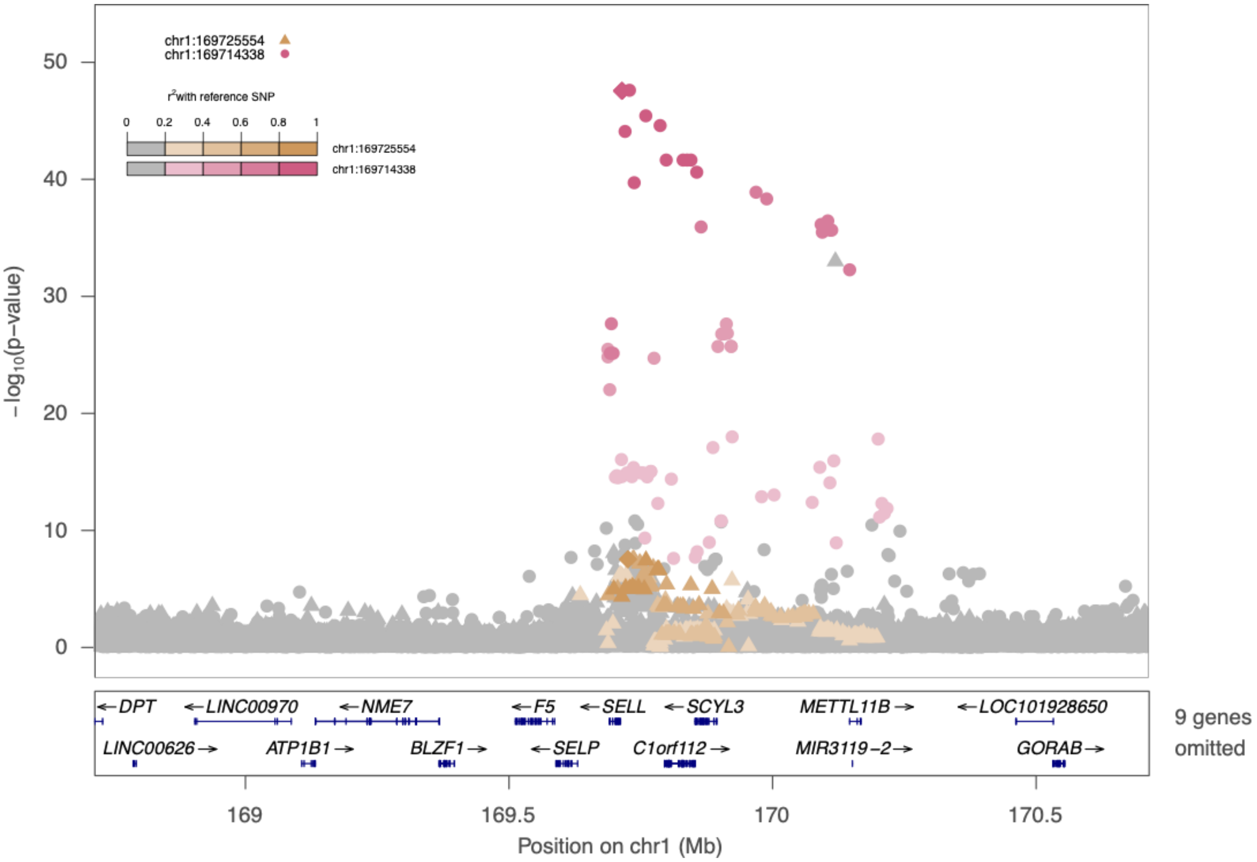


**Fig S13. Lead signal of E-selectin on *SELE*.** Lead signals are chr1:169714338_C_A (rs139825795) and chr1:169725554_C_T (rs3917434).

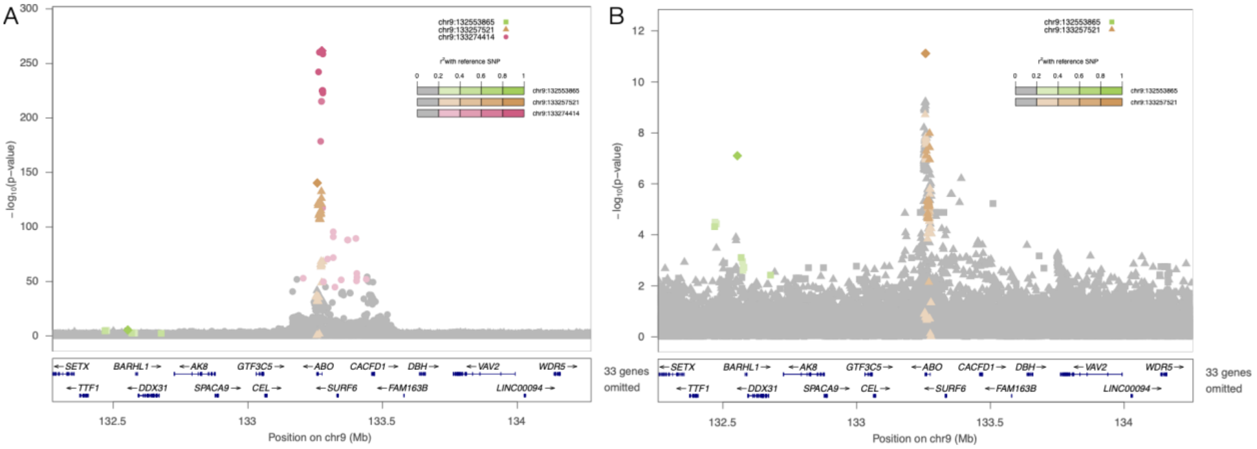


**Fig S14. Lead signal of E-selectin on *ABO*, prior to and after conditioning on previous literature reported variants. A.** Prior to conditional analysis, lead signals detected are chr9:133274414_G_A (rs947073006), chr9:133257521_TC_T (rs8176719), chr9:132553865_A_G (rs374594061). **B.** After conditional analysis, lead signals detected are chr9:133257521_TC_T (rs8176719), chr9:132553865_A_G (rs374594061).

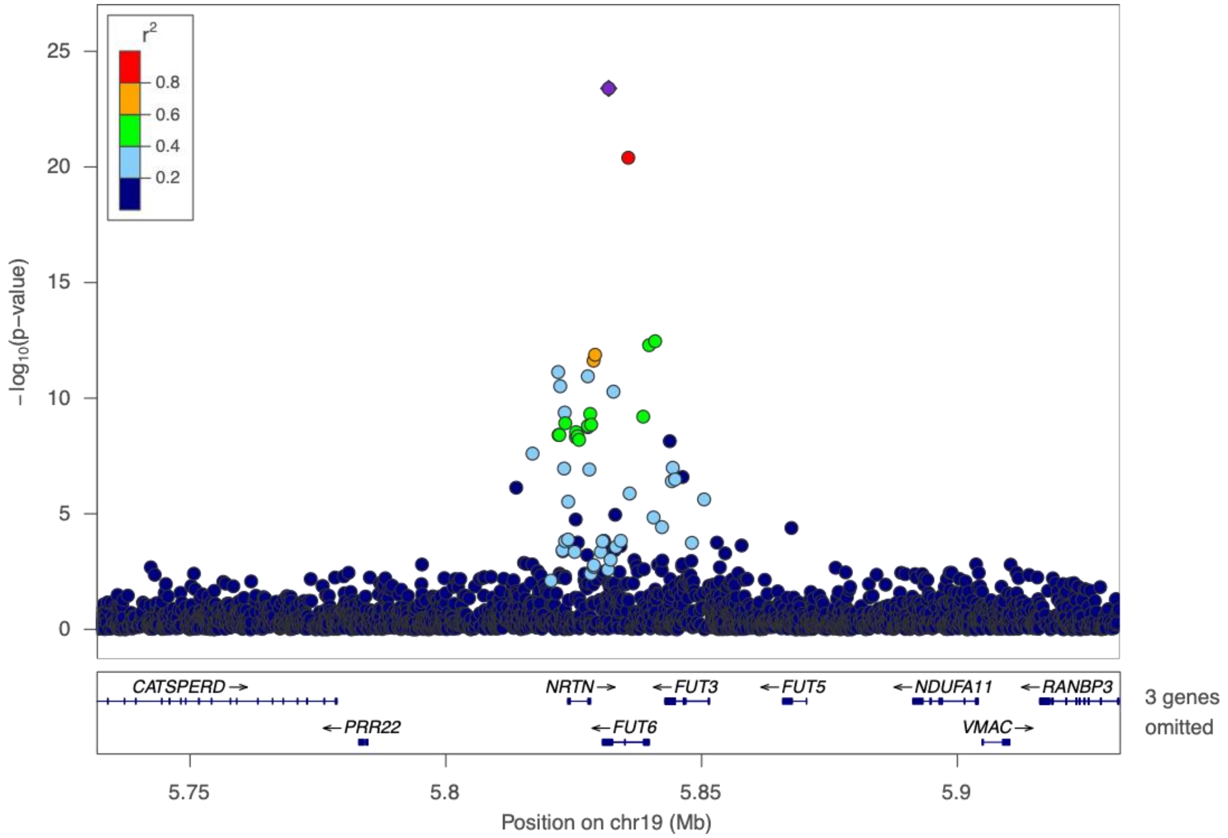


**Fig S15. Lead signal of E-selectin on *FUT6*.** Lead signal is chr19:5831829_T_C (rs17855739).

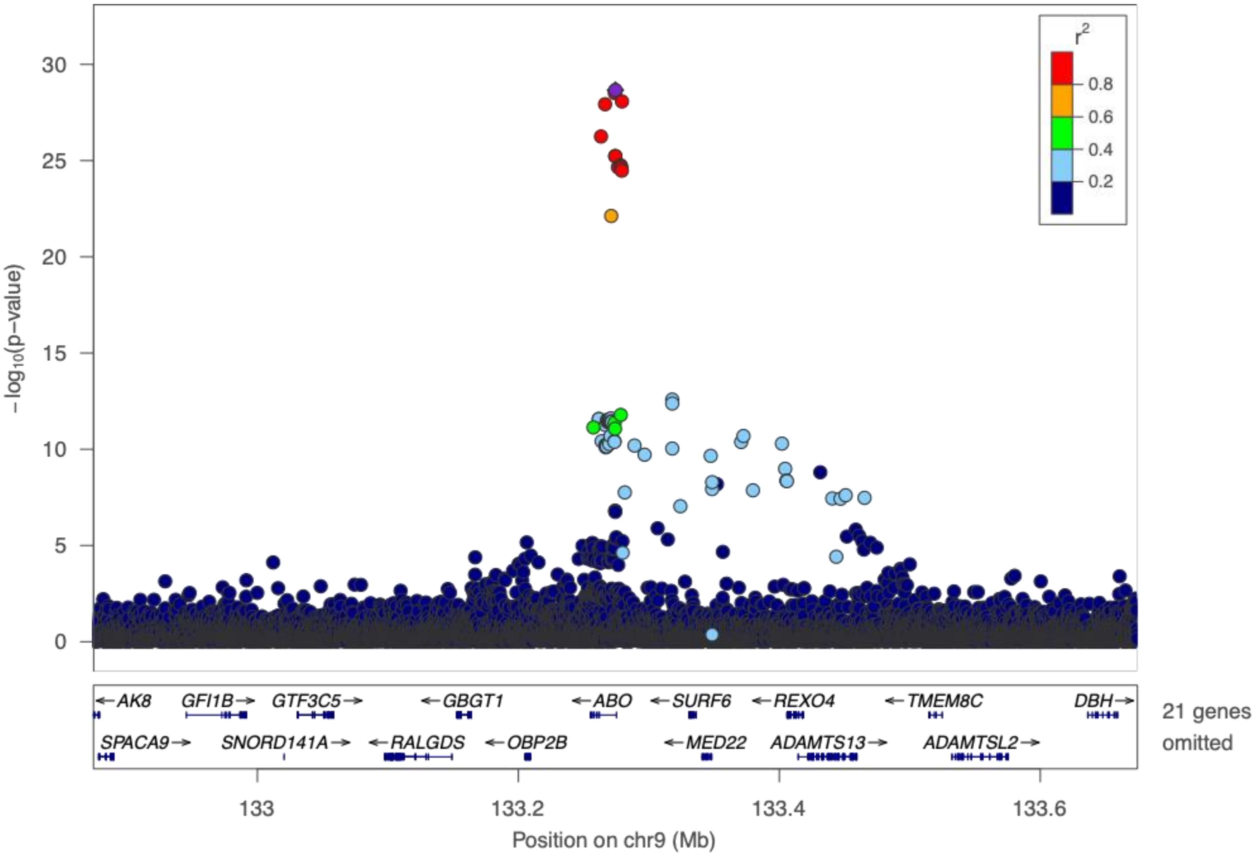


**Figure S16. Lead signal of ICAM-1 on *ABO*.** Lead signal is chr9:133274414_G_A (rs947073006).

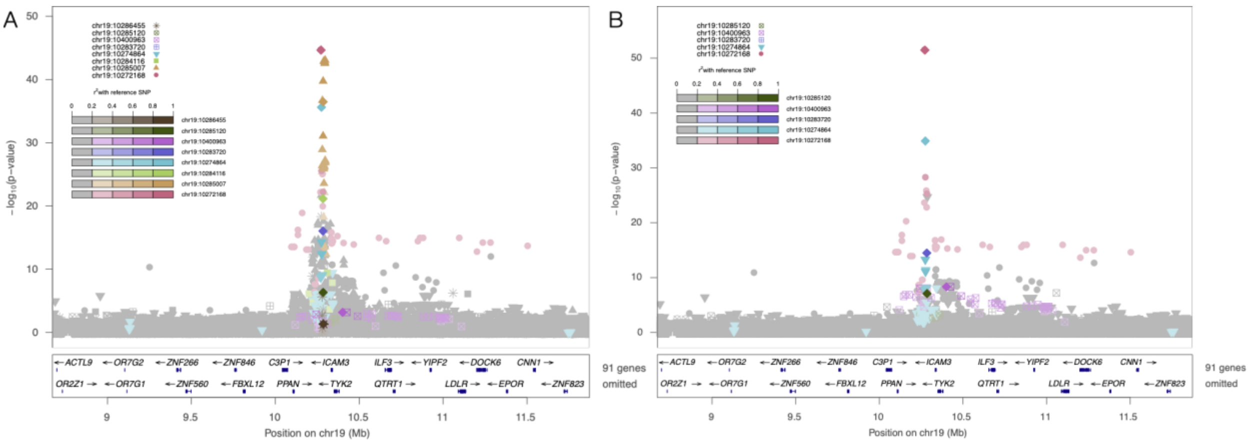


**Figure S17. Lead signals of ICAM-1 in *ICAM1*, prior to and after conditioning on previous literature reported variants. A.** Prior to conditional analysis, lead signals detected are chr19:10272168_G_C (rs11575071), chr19:10285007_G_A (rs5498), chr19:10284116_A_G (rs1799969), chr19:10274864_T_A (rs5491), chr19:10283720_C_G (rs139053442), chr19:10400963_G_T (rs28382777), chr19:10285120_T_C (rs5030400), chr19:10286455_A_G (rs281436). **B.** After conditional analysis, leads signals detected are chr19:10272168_G_C (rs11575071), chr19:10274864_T_A (rs5491), chr19:10283720_C_G (rs139053442), chr19:10400963_G_T (rs28382777), chr19:10285120_T_C (rs5030400).

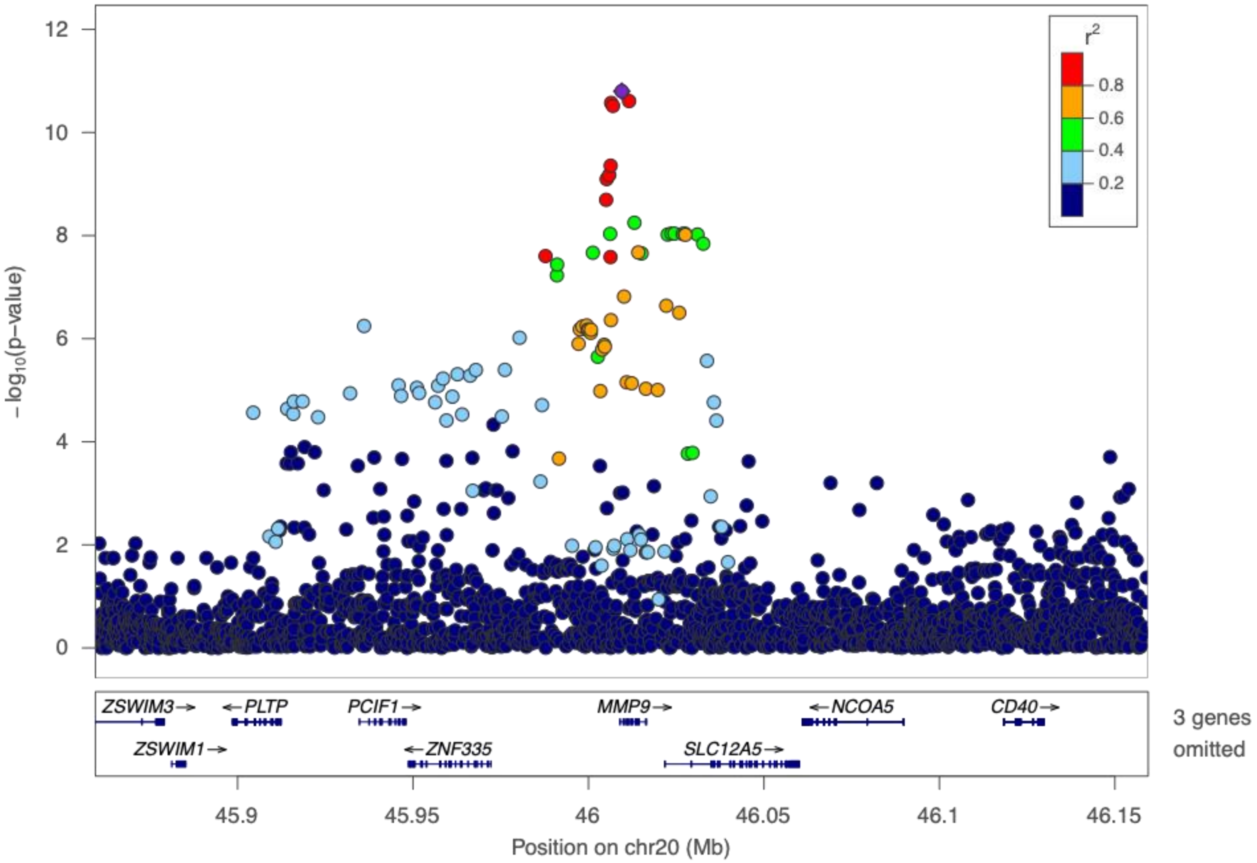


**Fig S18. Lead signals of MMP-9 on *MMP9*.** Lead signal is chr20:46009497_C_T (rs3918249).

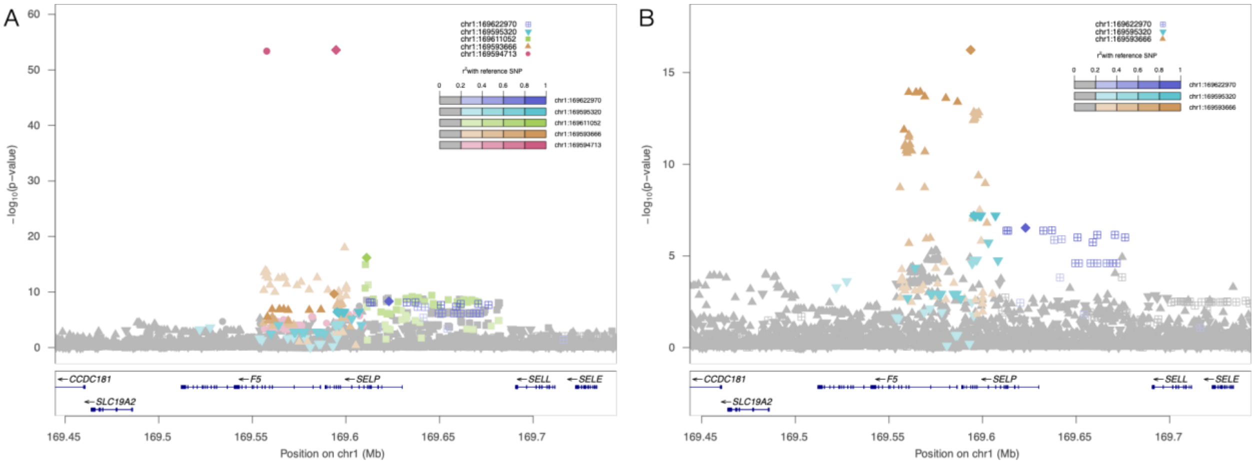


**Figure S19. Lead signals of P-selectin on *SELP*, prior to and after conditioning on previous literature reported variants. A.** Prior to conditional analysis, lead signals detected are chr1:169594713_G_T (rs6136), chr1:169593666_T_C (rs6128), chr1:169611052_T_C (rs2235302), chr1:169595320_G_A (rs3917825), chr1:169622970_C_A (rs3917677). **B.** After conditional analysis, lead signals detected are chr1:169593666_T_C (rs6128), chr1:169595320_G_A (rs3917825), chr1:169622970_C_A (rs3917677).

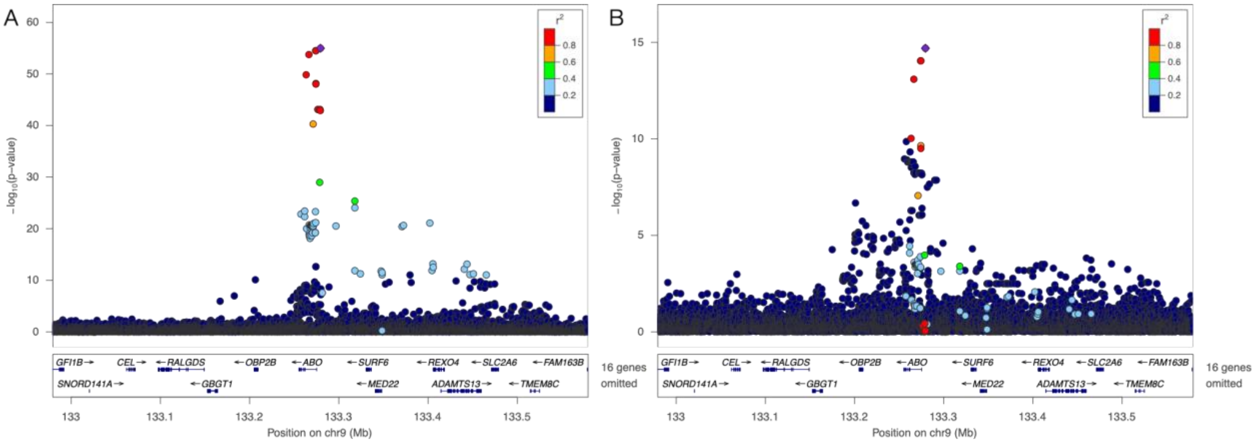


**Fig S20. Lead signal of P-selectin on *ABO*, prior to and after conditioning on previous literature reported variants. A.** Prior to conditional analysis, the lead signal detected is chr9:133279427_C_T (rs635634) ($p=1.0x10^{-55}$). **B.** After conditional analysis, the lead signal detected is still chr9:133279427_C_T (rs635634) while significance is attenuated ($p=2.0x10^{-15}$).

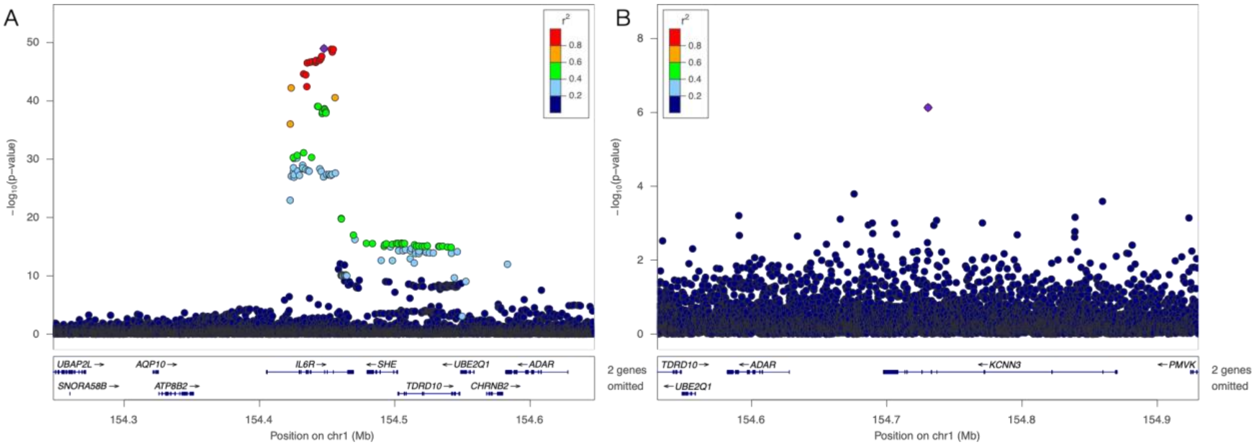


**Fig S21. Lead signals of IL-6 on *IL6R*, prior to and after conditioning on previous literature reported variants. A.** Prior to conditional analysis, the lead signal detected is chr1:154447611_A_G (rs61812598). **B.** After conditional analysis, the lead signal detected is chr1:154730517_T_C (rs568587329).

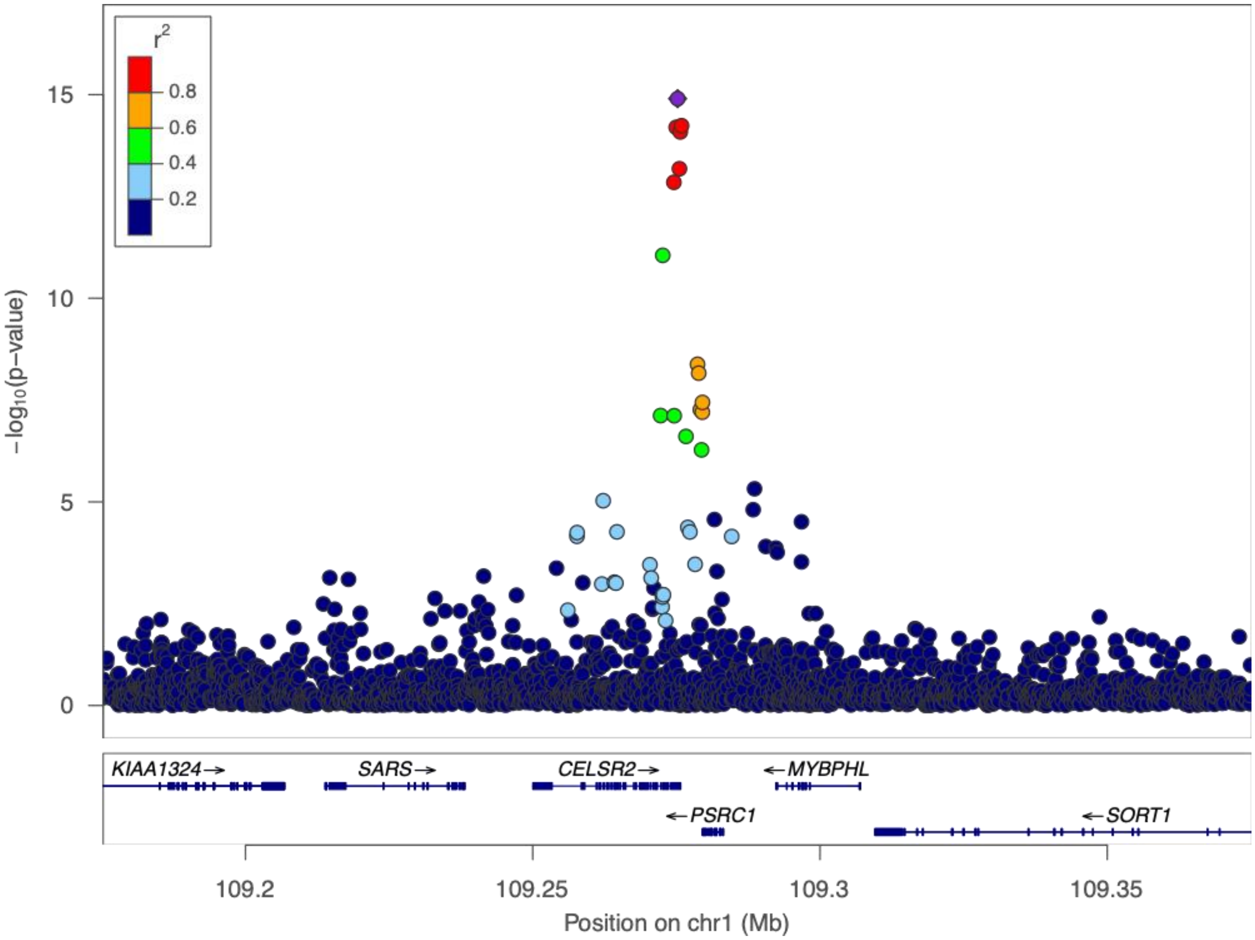


**Fig S22. Lead signals of Lp-PLA2 Activity on *CELSR2*.** Lead signal is chr1:109275216_C_T (rs660240).

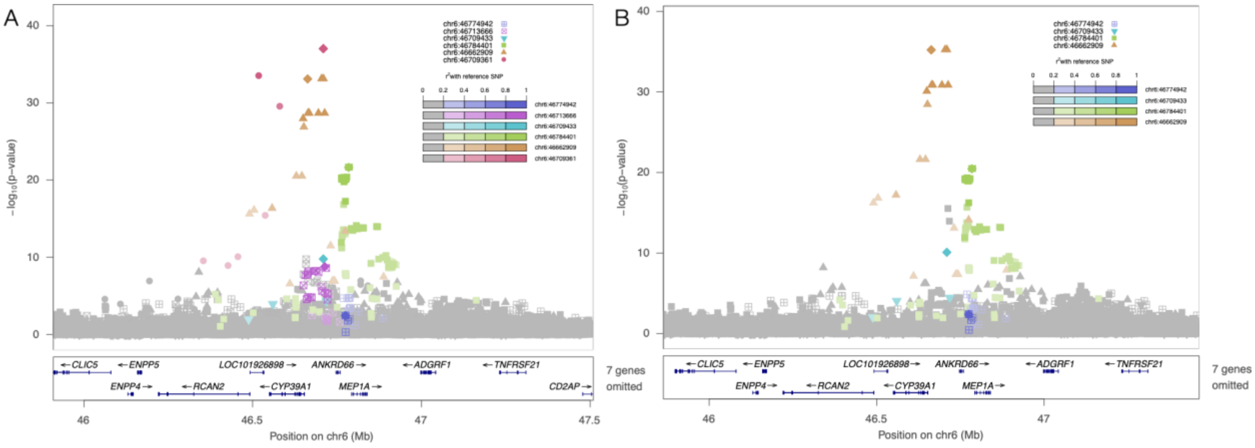


**Fig S23. Lead signals of Lp-PLA2 Activity on *PLA2G7*, prior to and after conditioning on previous literature reported variants. A.** Prior to conditional analysis, lead signals detected are chr6:46709361_A_C (rs76863441), chr6:46662909_G_T (rs144007943), chr6:46784401_A_G (rs74479543), chr6:46709433_G_A (rs144067869), chr6:46713666_G_A (rs1421372), chr6:46774942_A_C (rs150641786). **B.** After conditional analysis, lead signals detected are chr6:46662909_G_T (rs144007943), chr6:46784401_A_G (rs74479543), chr6:46709433_G_A (rs144067869), chr6:46774942_A_C (rs150641786).

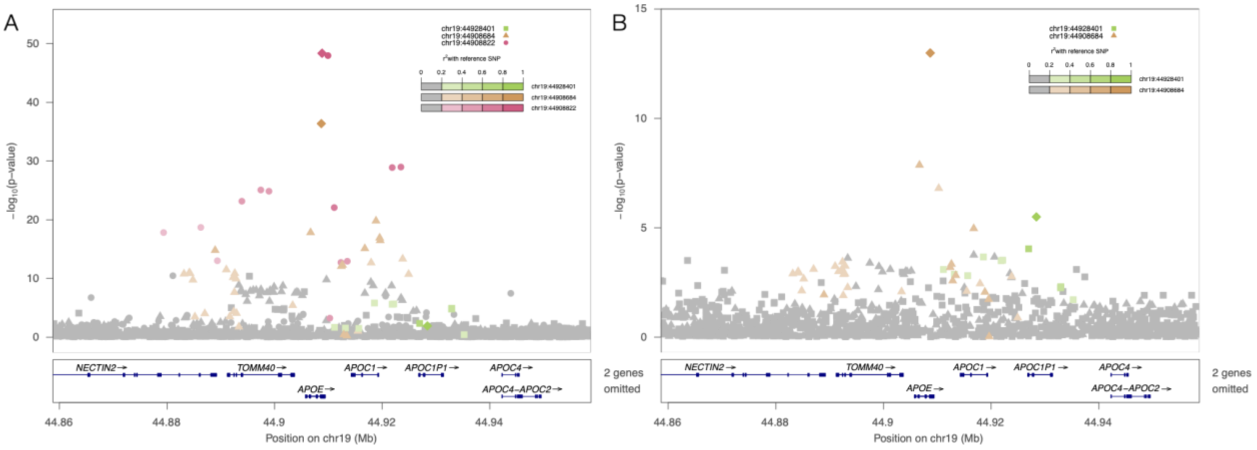


**Fig S24. Lead signals of Lp-PLA2 Activity on *APOE*, prior to and after conditioning on previous literature reported variants. A.** Prior to conditional analysis, lead signals detected are chr19:44908822_T_C (rs7412), chr19:44908684_C_T (rs429358), chr19:44928401_G_A (rs8106813). **B.** After conditional analysis, lead signals detected are chr19:44908684_C_T (rs429358), chr19:44928401_G_A (rs8106813).

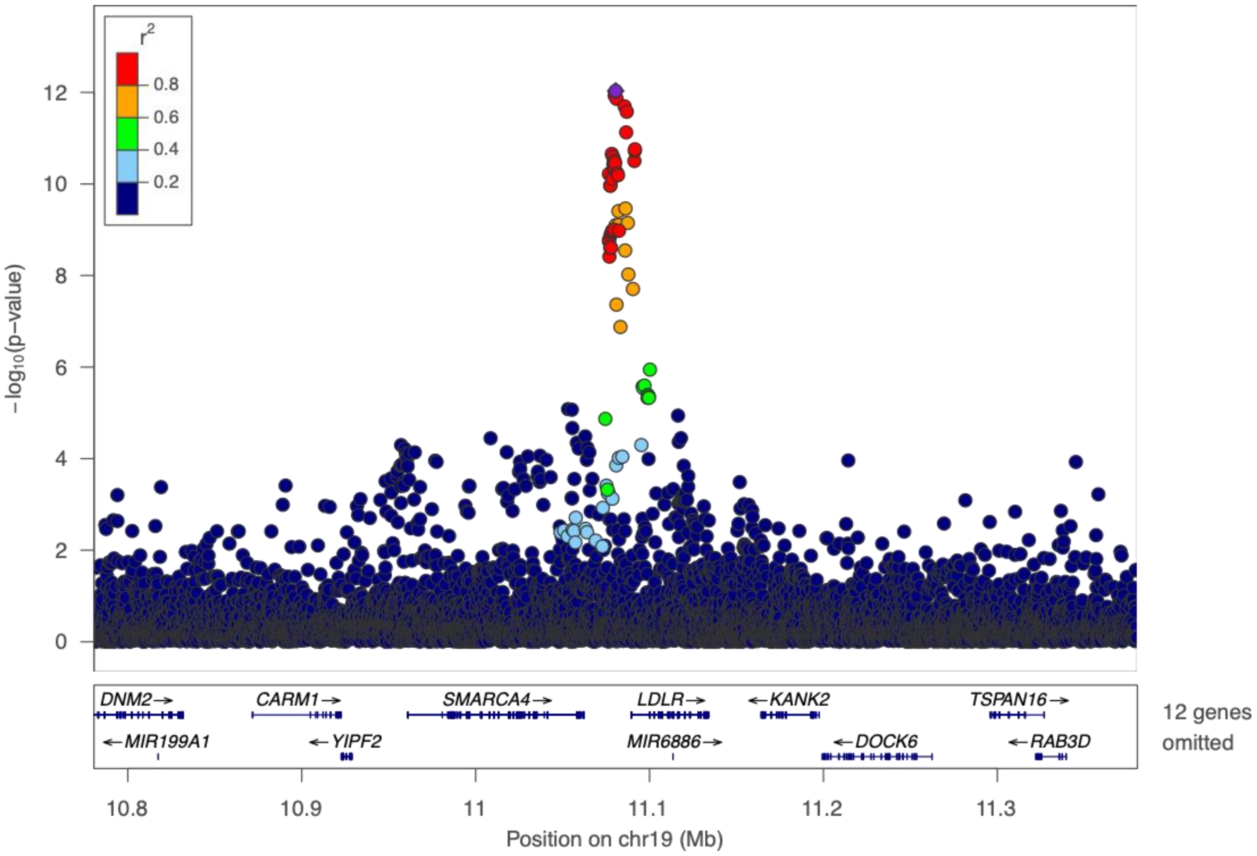

**Fig S25. Lead signals of Lp-PLA2 Activity on *LDLR*.** Lead signal is chr19:11080521_A_G (rs114846969).

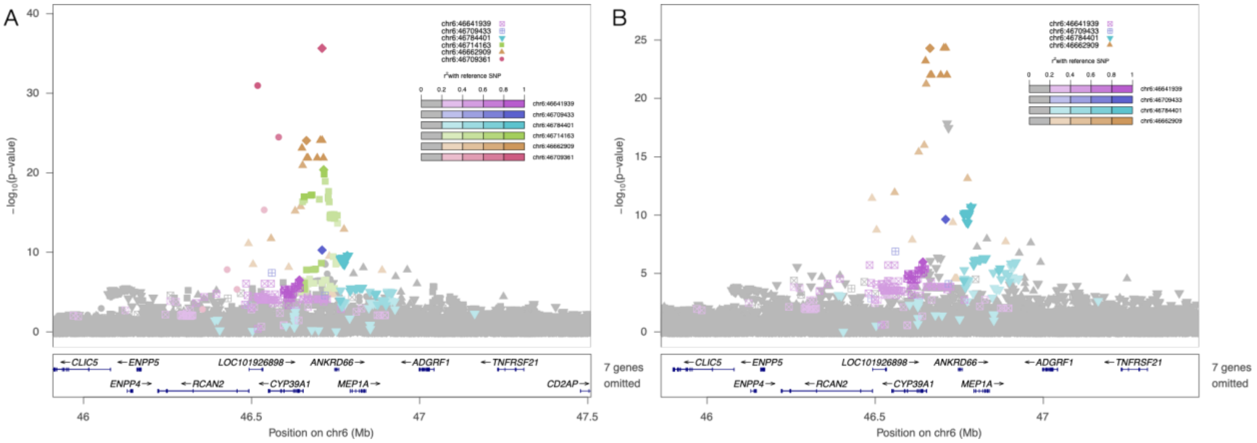

**Fig S26. Lead signals of Lp-PLA2 Mass on *PLA2G7*, prior to and after conditioning on previous literature reported variants. A.** Prior to conditional analysis, lead signals detected are chr6:46709361_A_C (rs76863441), chr6:46662909_G_T (rs144007943), chr6:46714163_G_T (rs6899519), chr6:46784401_A_G (rs74479543), chr6:46709433_G_A (rs144067869), chr6:46641939_C_T (rs73471140). **B.** After conditional analysis, lead signals detected are chr6:46662909_G_T (rs144007943), chr6:46784401_A_G (rs74479543), chr6:46709433_G_A (rs144067869), chr6:46641939_C_T (rs73471140).

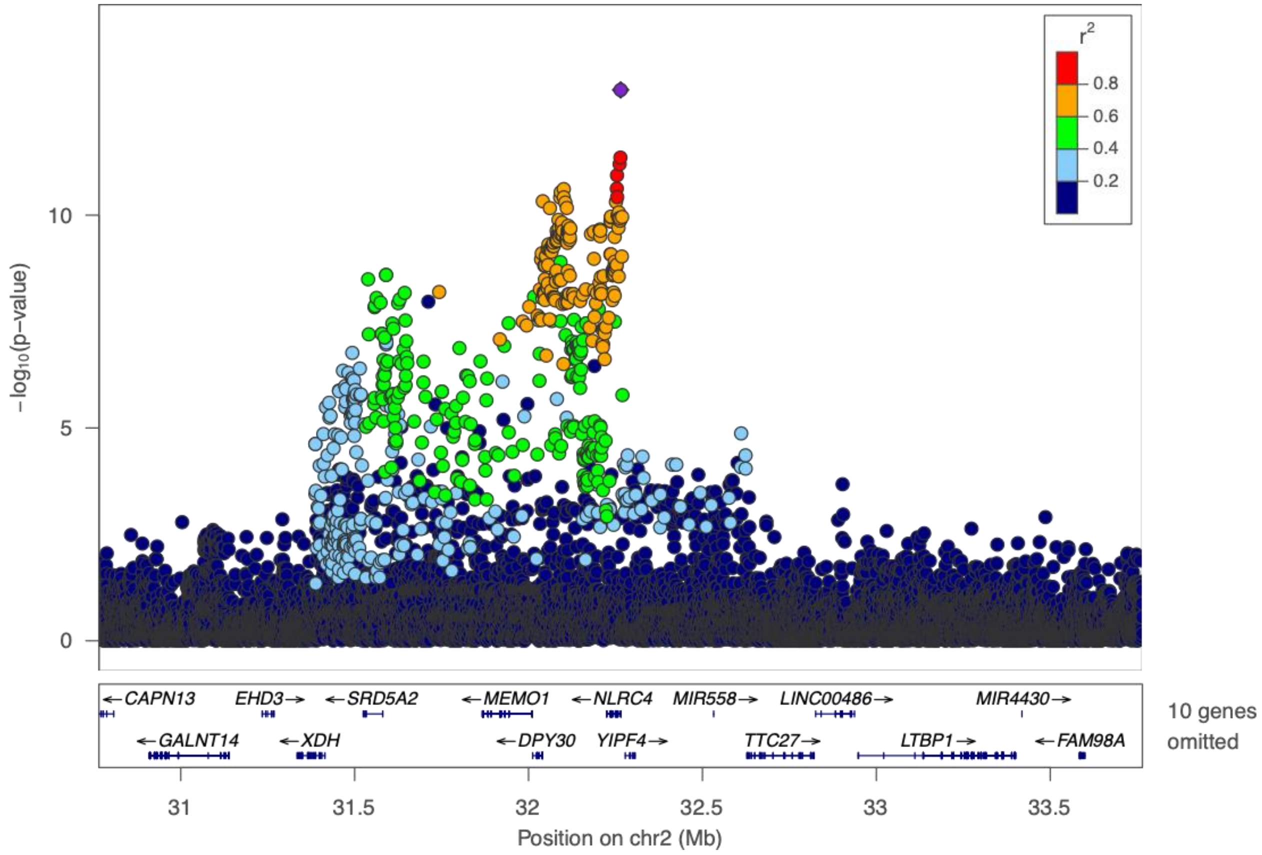


**Fig S27. Lead signals of IL-18 on *NLRC4*.** Lead signal is chr2:32264782_C_T (rs385076).

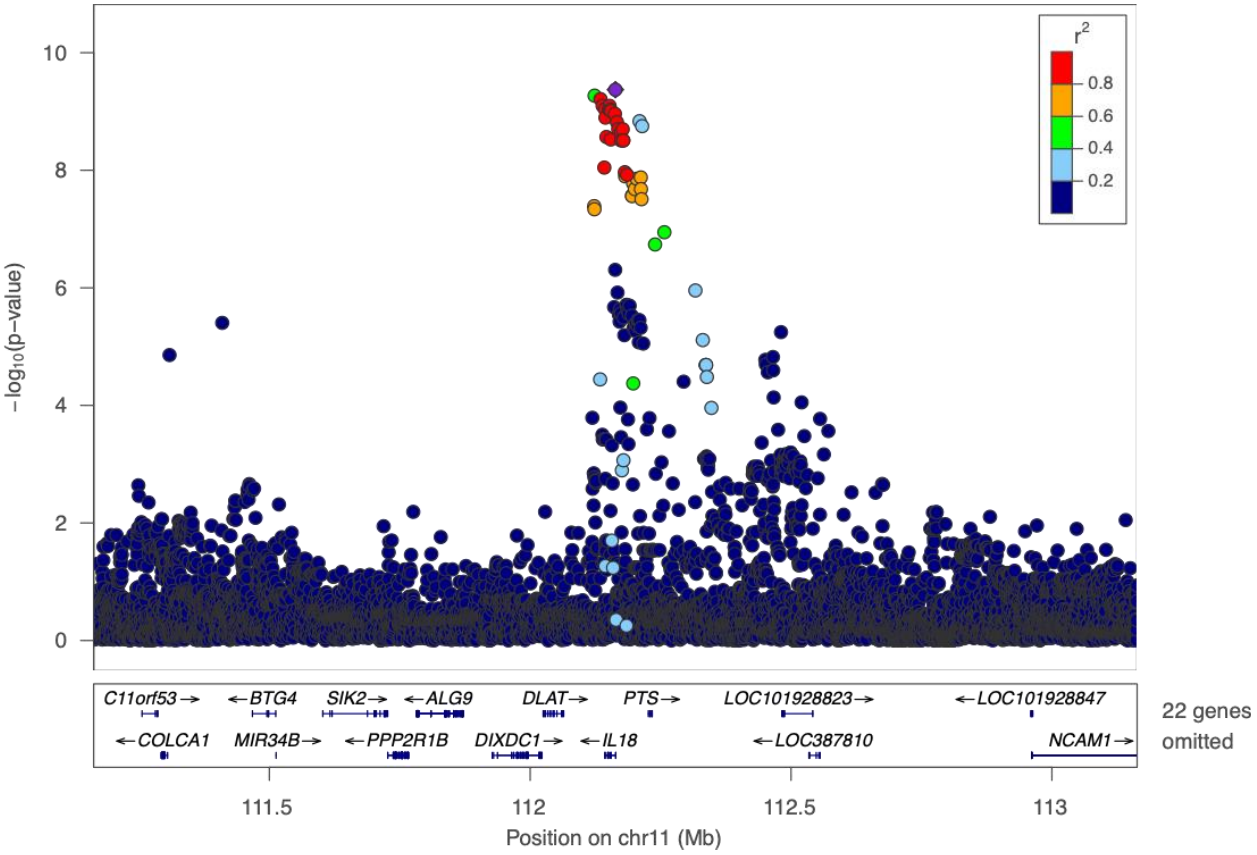


**Fig S28. Lead signals of IL-18 on *IL18*.** Lead signal is chr11:112163339_TA_T (rs397826848).

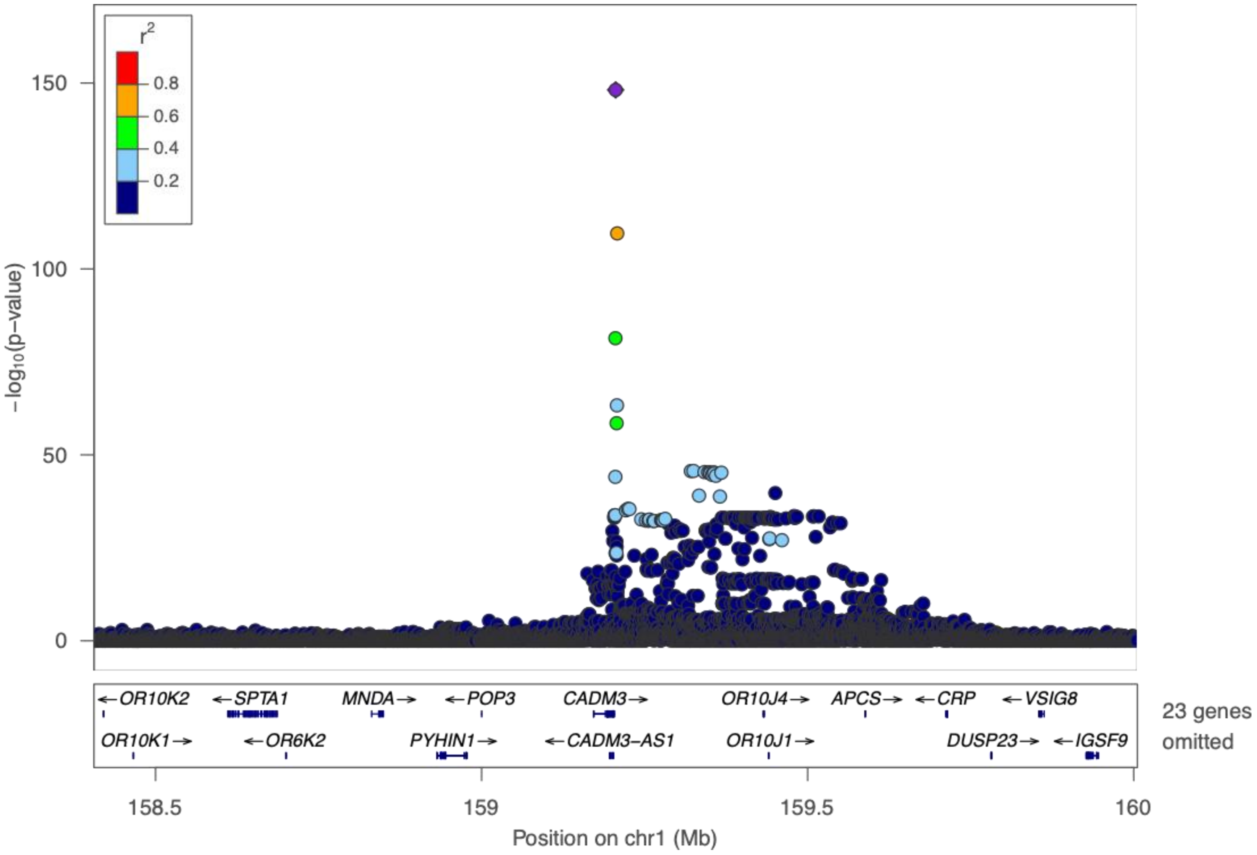


**Fig S29. Lead signal of MCP-1 on *ACKR1*.** Lead signal is chr1:159205564_A_G (rs12075).

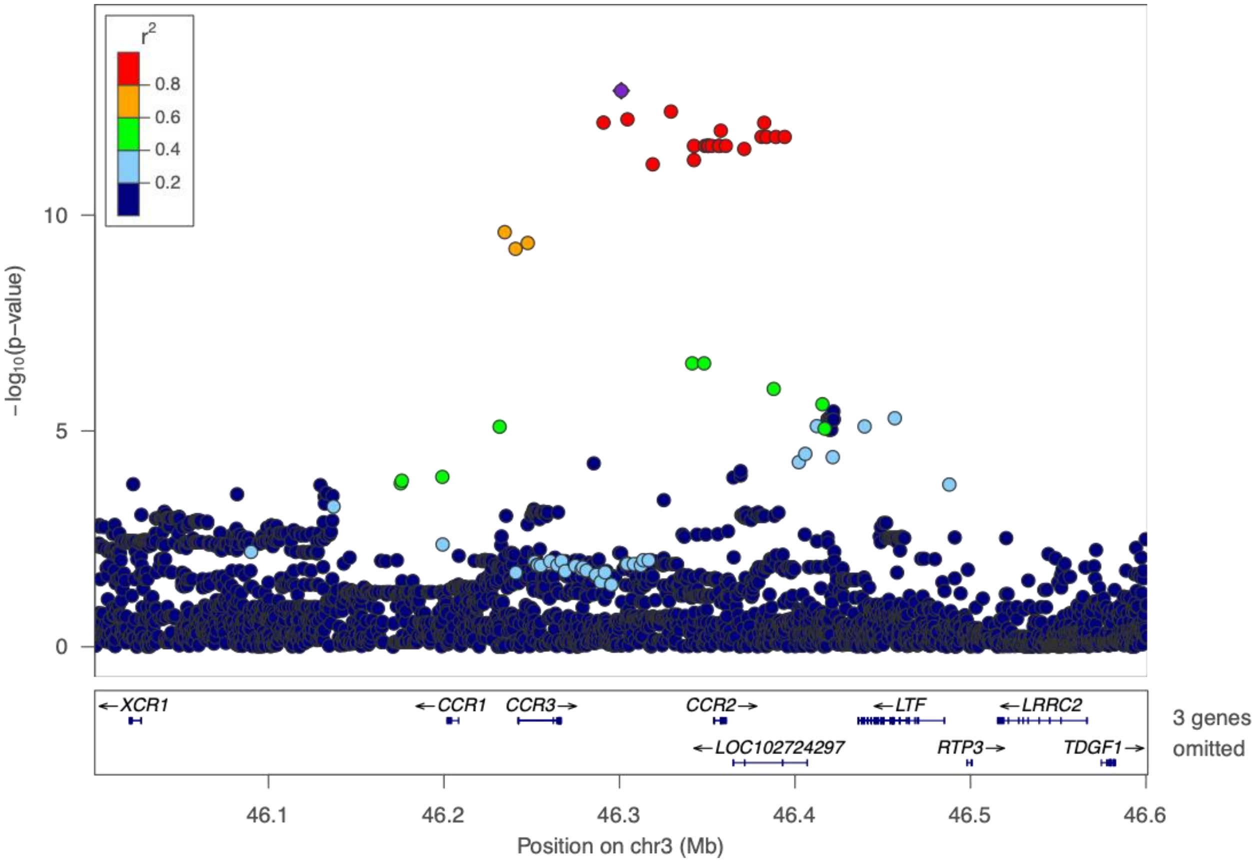


**Fig S30. Lead signal of MCP-1 on *CCR2*.** Lead signal is chr3:46301011_G_T (rs12495098).

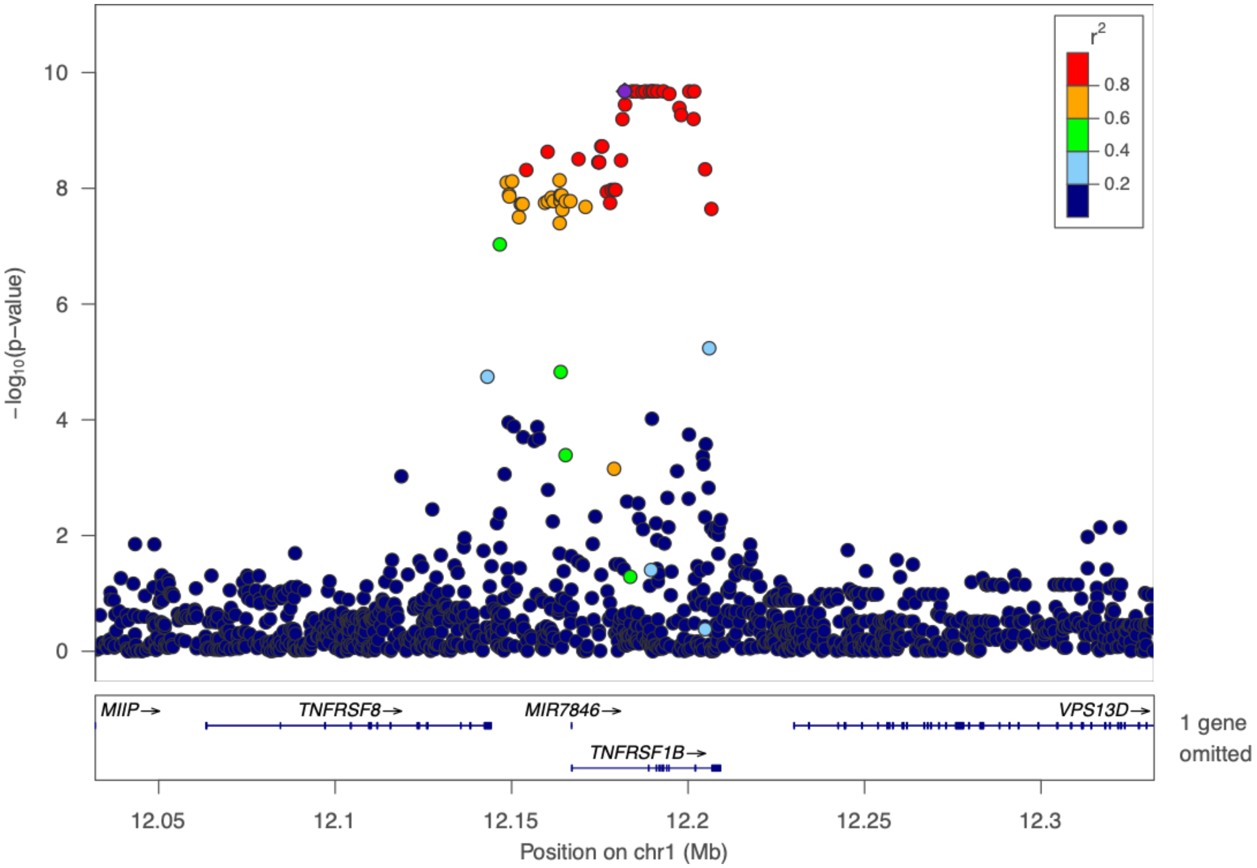


**Fig S31. Lead signal of TNFR2 on *TNFR2*.** Lead signal is chr1:12182039_T_C (rs519064).

<table>
<tr><td>A. CD40</td><td>B. CRP</td><td>C. E-selectin</td></tr>
<tr><td>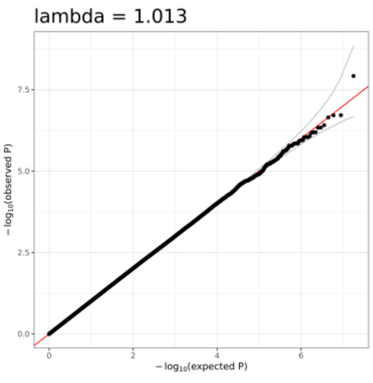</td><td>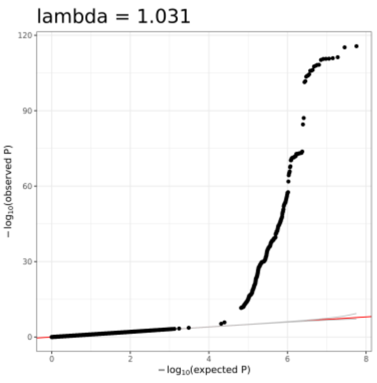</td><td>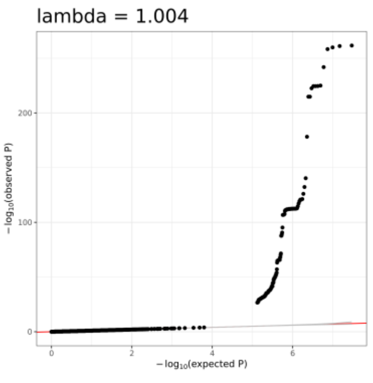</td></tr>
<tr><td>D. ICAM-1</td><td>E. IL1_beta</td><td>F. IL-6</td></tr>
</table>

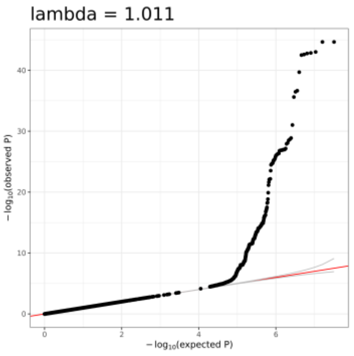


G. IL-8

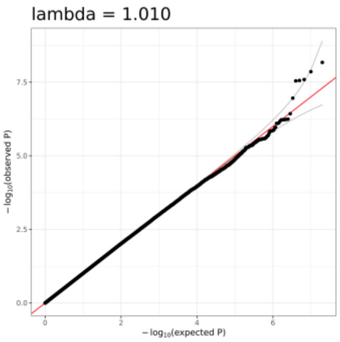


H. IL-10

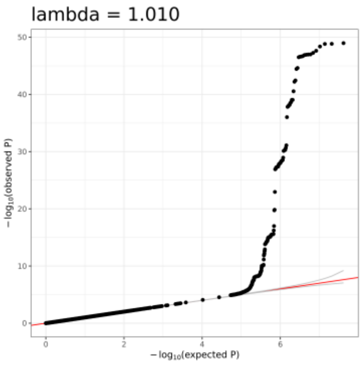


I. IL-18

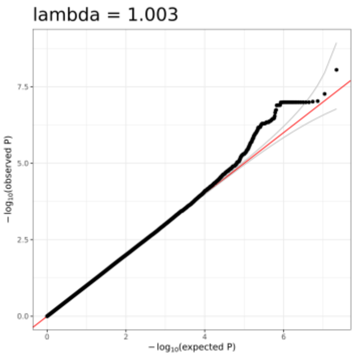


J. isoprostane-8-epi-pgf2a

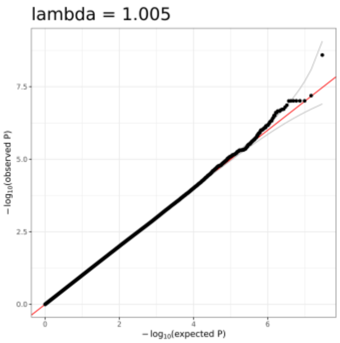


K. Lp-PLA2 Activity

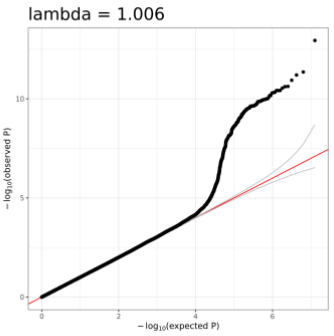


L. Lp-PLA2 Mass

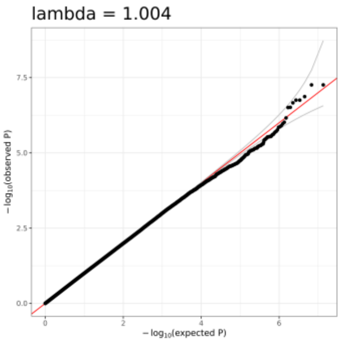


M. MCP-1

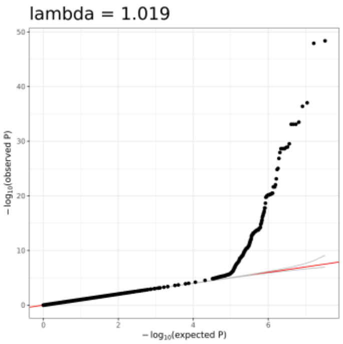


N. MMP-1

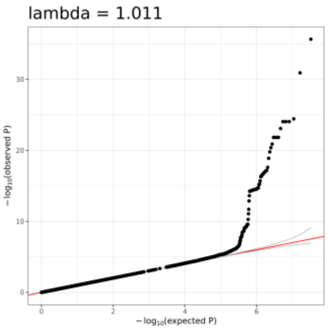


O. MMP-9

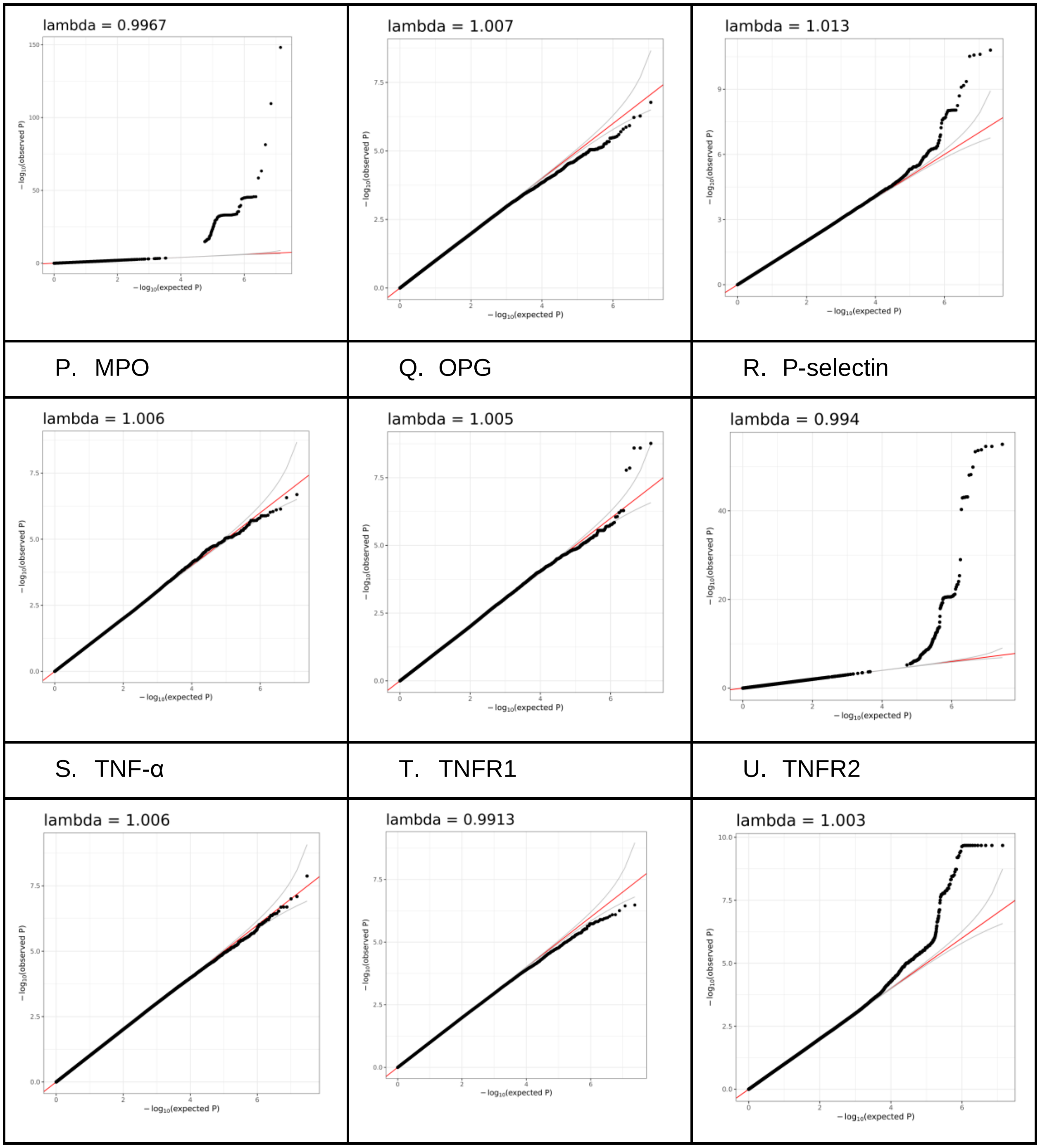


**Fig S32. QQ plots of marginal single variant association analysis of each trait.** A. Cluster of Differentiation 40 (CD40); B. C-Reactive Protein (CRP); C. E-selectin ; D. Intercellular Adhesion Molecule 1 (ICAM-1); E. Interleukin-10 (IL-10); F. Interleukin-18 (IL-18); G. Interleukin-1β (IL-1β); H. Interleukin-6 (IL-6); I. Interleukin-8 (IL-8); J. 8-iso Prostaglandin F2α (isoprostane-8-epi-pgf2α); K. Lipoprotein-associated phospholipase A2 (Lp-PLA2) Activity; L. Lipoprotein-associated phospholipase A2 (Lp-PLA2) Mass; M. Monocyte Chemoattractant Protein-1 (MCP-1); N. Matrix Metalloproteinase-1 (MMP-1); O. Matrix metalloproteinase-9 (MMP-9); P. Myeloperoxidase (MPO); Q. Osteoprotegerin (OPG); R. P-selectin; S. Tumor Necrosis Factor-α

(TNF-α); T. Tumor Necrosis Factor-α Receptor 1 (TNFR1); U. Tumor Necrosis Factor Receptor 2 (TNFR2).

#### Cohorts Acknowledgements

Participating cohorts have been described in prior TOPMed publications (1–3). Very brief descriptions, cohort specific acknowledgements, and cohort-specific methods for inflammation traits are listed in Tables S1 and S2.
